# Genealogical structure changes as range expansions transition from pushed to pulled

**DOI:** 10.1101/2020.12.29.424763

**Authors:** Gabriel Birzu, Oskar Hallatschek, Kirill S. Korolev

## Abstract

Range expansions accelerate evolution through multiple mechanisms including gene surfing and genetic drift. The inference and control of these evolutionary processes ultimately relies on the information contained in genealogical trees. Currently, there are two opposing views on how range expansions shape genealogies. In invasion biology, expansions are typically approximated by a series of population bottlenecks producing genealogies with only pairwise mergers between lineages—a process known as the Kingman coalescent. Conversely, traveling-wave models predict a coalescent with multiple mergers, known as the Bolthausen–Sznitman coalescent. Here, we unify these two approaches and show that expansions can generate an entire spectrum of coalescent topologies. Specifically, we show that tree topology is controlled by growth dynamics at the front and exhibits large differences between pulled and pushed expansions. These differences are explained by the fluctuations in the total number of descendants left by the early founders. High growth cooperativity leads to a narrow distribution of reproductive values and the Kingman coalescent. Conversely, low growth cooperativity results in a broad distribution, whose exponent controls the merger sizes in the genealogies. These broad distribution and non-Kingman tree topologies emerge due to the fluctuations in the front shape and position and do not occur in quasi-deterministic simulations. Overall, our results show that range expansions provide a robust mechanism for generating different types of multiple mergers, which could be similar those observed in populations with strong selection or high fecundity. Thus, caution should be exercised in making inferences about the origin of non-Kingman genealogies.

**Significance statement:** Spatial dynamics are important for understanding genetic diversity in many contexts, such as cancer and infectious diseases. Coalescent theory offers a powerful framework for interpreting and predicting patters of genetic diversity in populations, but incorporating spatial structure into the theory has proven difficult. Here, we address this long-standing problem by studying the coalescent in a spatially expanding population. We find the topology of the coalescent changes depending on the growth dynamics at the front. Using analytical arguments, we show that the transition between coalescent topologies is universal and is controlled by a parameter related to the expansion velocity. Our theory makes precise predictions about the effects of population dynamics on genetic diversity at the expansion front, which we confirm in simulations.

## Introduction

The genealogy of a population provides a window into its past dynamics and future evolution. By analyzing the relative lengths of different branches in the genealogical tree, we can estimate mutation rates and the strength of genetic drift [1], or infer historical population sizes [2] and patterns of genetic exchange between species [3]. At the same time, we can use the structure of genealogies to make predictions about the speed of evolution [4] and even answer important practical questions, such as what the next strain of influenza will be [5].

Typically, the full ancestry of the population is not known and has to be inferred from DNA samples using theoretical models. The most widely-used model is the Kingman coalescent [6,7]. The Kingman coalescent describes the genealogies of a well-mixed population of constant size, in which all mutations are neutral. Because of its simplicity, many statistical properties of the Kingman coalescent can be calculated exactly [7]. These mathematical results have formed the basis of many commonly-used techniques to infer genealogical trees from DNA sequences. The defining characteristics of the trees generated from the Kingman coalescent are a large number of early mergers and long branches close to the common ancestor. Importantly, the Kingman coalescent contains only pairwise mergers between lineages. However, several studies have attempted to test these predictions directly in real populations and found significant deviations [8–11].

To resolve the inconsistencies between observed genetic diversity and theoretical predictions, numerous extensions of the classic Kingman coalescent have been proposed [12–16]. For example, many studies have analyzed the effects of time-dependent population sizes and spatial structure on the coalescent [2, 17]. Despite providing better fits to the data, this generalized Kingman coalescent does not capture some of the qualitative features of empirical genealogies—namely the existence of multiple mergers in the genealogical trees [18, 19].

Over time, several mechanisms that give rise to coalescents with multiple mergers have also been proposed. Theoretical studies have shown that highly fecund populations have multiple mergers in their genealogies [20,21]. Selective sweeps can also lead to fat-tailed distributions in the number of offspring. Mathematically, the genealogies of such populations can be described by a more general coalescent model known as the Λ-coalescent [20,22]. However these mechanisms have limited applicability—most species have few offspring and typical population sizes and selective pressures are unlikely to have a large effect on genealogies [23–25]. Here, we show that a ubiquitous demographic mechanism generates genealogical trees with a wide range of topologies, including topologies with exclusively pairwise mergers as well as topologies with multiple mergers. This mechanism relies on unusually large genetic drift at the leading edge of expanding population fronts. Such expansions can occur in a variety of contexts, such as range expansions [26], range shifts due to climate change [27], or the growth of bacterial colonies [28, 29] and tumors [30, 31].

Despite their importance, very little is known about the genealogies of spatially expanding populations. Two approaches have been used previously to study this problem, often leading to very different conclusions [32–34]. The most common approach is to approximate spatial expansions by a series of discrete bottlenecks at the front [23, 32, 35]. This is known as the serial bottleneck approximation and it implicitly assumes that genealogies along the expansion are described by a series of replacement events (as illustrated in Fig. 1a, c), while those at the leading edge are described by the Kingman coalescent, with a potentially time-dependent population size [33, 36]. The Kingman structure of genealogies has also been recently proven for a certain class of range expansions with negative growth rates at the leading edge [37]. An alternative approach, introduced in Ref. [34], is based on an analogy between spatial expansions and traveling waves describing the increase in fitness in a population of constant size under strong selection [38–40]. Using heuristic arguments, supported by extensive numerical simulations, Brunet et al. conjectured that expansions under the Fisher-Kolmogorov-Petrovsky-Piskunov (FKPP) universality class are described by a different type of coalescent, known as the Bolthausen-Sznitman coalescent^1^ [34]. Unlike the standard Kingman coalescent, in which only pairwise mergers between branches are allowed, the Bolthausen-Sznitman coalescent is characterized by large merger events, during which a finite fraction of branches can coalesce simultaneously [42, 43]. Despite subsequent investigations, reconciling these two diametrically opposed points of view is still an open problem [33, 36, 40].

**Figure 1:**
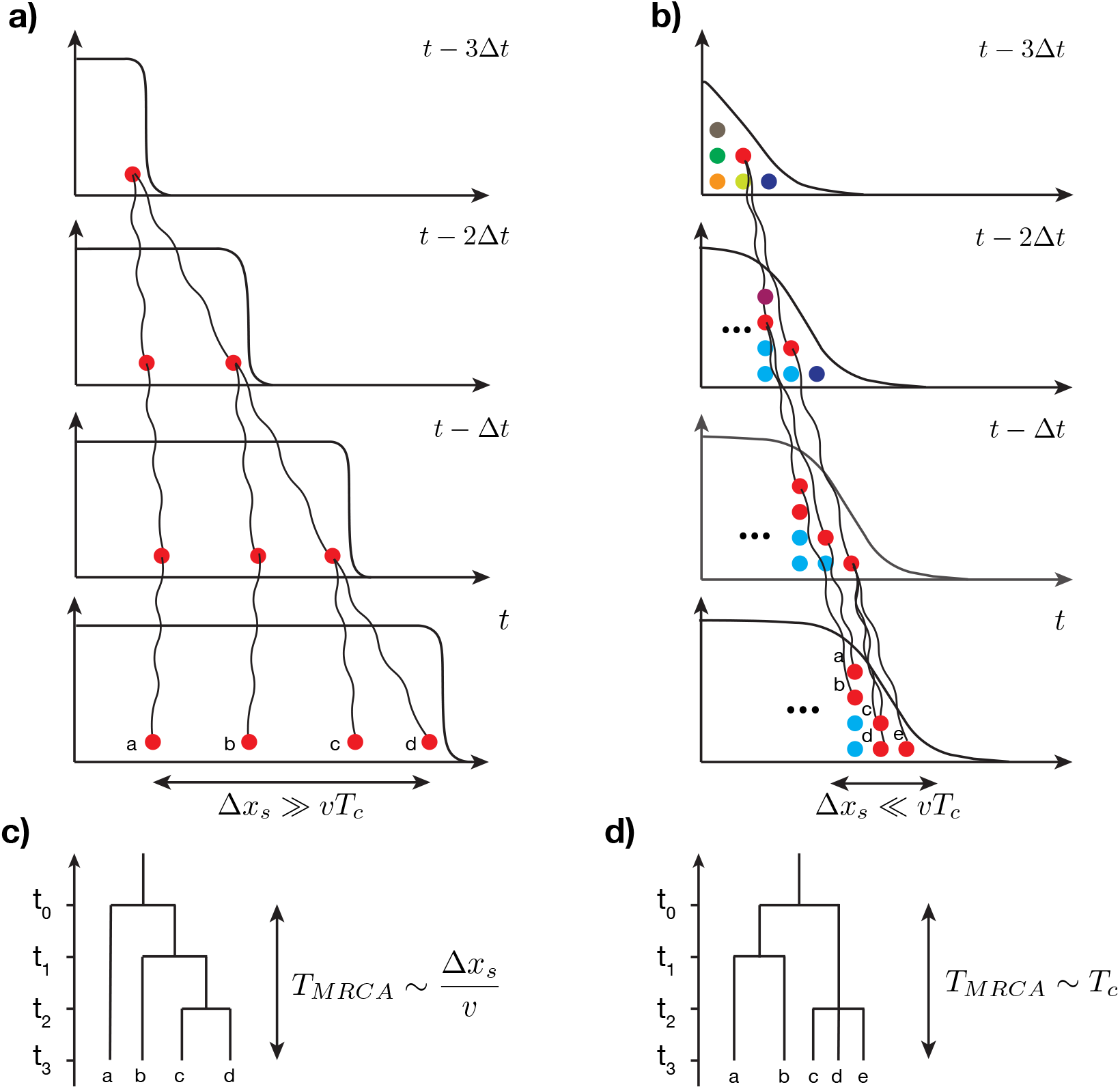
Shape of genealogies in expanding populations depends on spatial location of sampled individuals. The genealogies in two limiting sampling regimes are shown schematically. **(a)** When sampling is done over large distances along the expansion, the coalescence time is mainly determined by the motion of the front. **(b)** In this regime, the lineage coalescence depends on spatial locations and genealogies correspond to a series of replacement events. **(c)** When sampling is done at the front, lineage coalescence is independent of spatial location and the motion of the front does not play an important role. **(d)** In this regime, a characteristic coalescence time *T_c_* emerges which is determined by the topology of the genealogical tree.

Recent studies by the authors point to a potential resolution of the above-mentioned contradiction [44,45]. Specifically, we examined whether population dynamics at the front could lead to differences in the rate of diversity loss during range expansions. Surprisingly, we found that density dependence in either growth or migration has large effects on genetic diversity. These effects can be grouped into three distinct regimes. When density dependence is positive and large—such as when growth and migration are highly cooperative, for example—the time scale over which diversity is lost scales linearly with the carrying capacity. This is the scaling expected from the Kingman coalescent and is consistent with the serial bottlenecks view. However, when cooperation is reduced, large fluctuations in density at the front tip lead to sublinear scaling, as would be expected if multiple mergers were present in the genealogies [7]. Finally, when cooperation is absent, the timescale of diversity loss scales logarithmically with the carrying capacity, as would be expected from a population described by the Bolthausen-Sznitman coalescent [7]. These results lead to a natural hypothesis, that these changes in the rate of diversity loss are a result of changes in the underlying genealogies, driven by large fluctuations in the low-density region of the front.

In this paper, we elucidate the connection between population dynamics and genealogies during expansion. We focus on understanding the topology of genealogies in the well-mixed region close to the front of the expansion (Fig. 1b, d). Using simulations, we obtain genealogical trees and examine how they change as growth dynamics vary. We indeed find that changes in growth cooperativity lead to a transition from the Kingman to a non-Kingman coalescent with multiple mergers. The fluctuations in the position and shape of the expansion front are crucial to these results because we observe only the Kingman coalescent when demographic fluctuations of the front are artificially suppressed.

To explain our findings, we developed an effective model of the expansion front using analytical arguments. We showed that the front can be treated as a well-mixed population with a broad distribution of number of offspring (reproductive values). The tail of the distribution follows a power law with an exponent that depends only on the ratio of the expansion velocity and the geometric mean of the growth and dispersal rates at low population densities. The topology of the genealogies is described by a Λ-coalescent and is in turn determined by the exponent [21, 46, 47]. Thus, the distribution of merger sizes in the genealogies of expanding populations is dependent on the growth dynamics.

## Simulation results

### Expansion model

We simulated a population expansion using a setup similar to the classic stepping stone model [48]. Specifically, we consider a one-dimensional landscape of demes (patches). For computational efficiency, we use a simulation box of *L* = 300 demes, which moves with the expansion front such that the box is approximately half-filled at all times. Each generation, individuals migrate between neighboring demes with probability *m*/2 and reproduce. The number of descendants is determined by the growth function that depends on the local population density (see Methods for details). On average, the population density increases to a maximum value set by the carrying capacity *N*. All individuals are resampled every generation, so demes that are at carrying capacity still experience genetic drift. As a result, the model reduces to a Wright-Fisher process in the bulk and a branching process with Poisson distributed number of offspring at the front.

### Methods

The detailed implementation of the sampling of descendants can be found in the SI, Sec. IV. For our purposes here, the change in the local population size *n_k_*, can be represented by a growth function *r*(*n_k_*), given by the following expression:

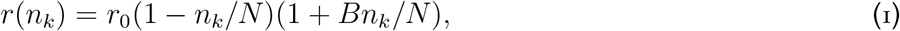

where *k* is the deme index and *r*_0_ is the growth rate at zero density. For convenience, we set the generation time to one and omit it from future expressions. The parameter *B* in (1) sets the growth cooperativity in the population. For *B* = 0, (1) is the widely-used logistic growth function [49,50], which has the maximum growth rate *r*(*n_k_*) = *r*_0_ at *n_k_* = 0. For *B* > 1, the position of the maximum shifts to *n_k_* > 0, and *r*(*n_k_*) becomes larger as *B* increases.

We showed previously that *B* in (1) controls the scaling between the carrying capacity *N* and the effective population size of the front *N_e_*, which we define as the time scale over which genetic diversity is lost. This dependence of *N_e_* on *N* changes from a linear function for *B* ≥ 4, to a power law for 2 < *B* < 4, and then to ln^3^ *N* for *B* < 2 [44]. We refer to the three expansion classes as fully pushed, semi-pushed, and pulled, respectively [44, 45]. This terminology reflects the fact that growth in pulled expansions occurs mainly at the edge of the front while, in semi-pushed and fully pushed expansions, it is in the bulk. We performed simulations with one value of B for each regime: *B* =10 for fully pushed expansions, *B* = 3.33 for semi-pushed expansions, and *B* = 0 for pulled expansions. Although our simulations are based on the specific growth and migration model detailed above, our theoretical results are model-independent (see below). Therefore, we do not expect any of our conclusions to change if different growth or migration models are used.

Genealogies can be obtained by storing all ancestral relationships. This approach, however, would severely constrain the population size and duration of our simulations. Instead, we keep track of genealogies by periodically assigning a unique label to every individual in the population. After assignment, the size of surviving clones—defined as a group of individuals with the same label—increases, while other clones become extinct. After a fixed number of generations Δ*t*, we relabel all individuals and store their previous labels. One can then trace the ancestry backward in time with temporal resolution Δ*t*. As long as Δ*t* is not too large compared to the generation time and the maximum clone size is small compared to the total population size, this procedure introduces only minor information loses in the genealogies for sample sizes much smaller than the carrying capacity.

### Descendant distribution in deterministic fronts

Without demographic fluctuations, the front profile *n_d_*(*ζ*) assumes a steady-state solution with a cutoff in the density determined by *n_d_*(*ζ_c_*) = 1, since the number of individuals cannot be less than one. Thus, for values of *ζ* > *ζ_c_*, the population density is zero. This density cutoff implies a maximum number of descendants^2^ *W_c_*, which can be calculated as discussed in the SI, Sec. I. Viewed backward in time, the ratio 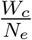 is the maximum fraction of lineages that can merge at the same time, where *N_e_* is the size of population at the front with a non-negligible probability of fixation. We find that 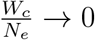 in the limit of large *N* (SI, Sec. I). Hence, pairwise mergers should dominate, leading to the Kingman coalescent.

### Spatial self-averaging

Range expansions are inherently heterogeneous in time and space. Therefore, ancestral relationships can in general depend on the times and locations of samples from the population. Consider two extreme sampling scenarios of either sampling individuals uniformly from the colonized range (Fig. 1a), or sampling all individuals from the front (Fig. 1b). In both cases, coalescent events primarily occur when ancestral lineages are at the front because genetic drift in the population bulk is much weaker. When two samples are taken from distant spatial locations, their lineages need to “wait” until both lineages are at the front. Viewed backward in time, this occurs when the front recedes past the left-most lineage (see Fig. 1a, c). Thus, in this sampling protocol, the shape of the genealogical tree explicitly depends on the spatial separation between the sampling locations. In contrast, there is no position-dependence when all individuals are sampled at the front because all lineages start merging at the same time (Fig. 1b, d).

Previous work suggests that lineages sampled at the front can be viewed as if they are part of a well-mixed population comoving with the front [44,51]. This approximation is valid on time scales longer than the mixing time.

To test if the mixing time *τ_m_* is indeed much shorter than the coalescence time, we tracked the spatial distribution of ancestors of individuals at the front. Specifically, we performed 30 independent simulations and sampled individuals from two spatial locations, one closer to the front and the other closer to the bulk. The inset in Fig. 2a shows the two sampling locations (blue and orange dots) together with the final front (grey line) for each run. The main panels show the distribution of ancestors of individuals from the two sampling locations shortly before the sampling time (Fig. 2a), and at a time close to *τ_m_* (Fig. 2b). Importantly, we found that the time necessary for the ancestor distributions to become independent of sampling location was much shorter than the time to reach the common ancestor for the whole front. For example, from Fig. 2b we estimated *τ_m_* ≈ 10^2^ generations, compared to *T_c_* ≈ 10^3^. These results show that the sampling positions do not affect genealogies and, therefore, the lineages can be considered *exchangeable,* which is a key requirement for describing them using the coalescent theory.

**Figure 2:**
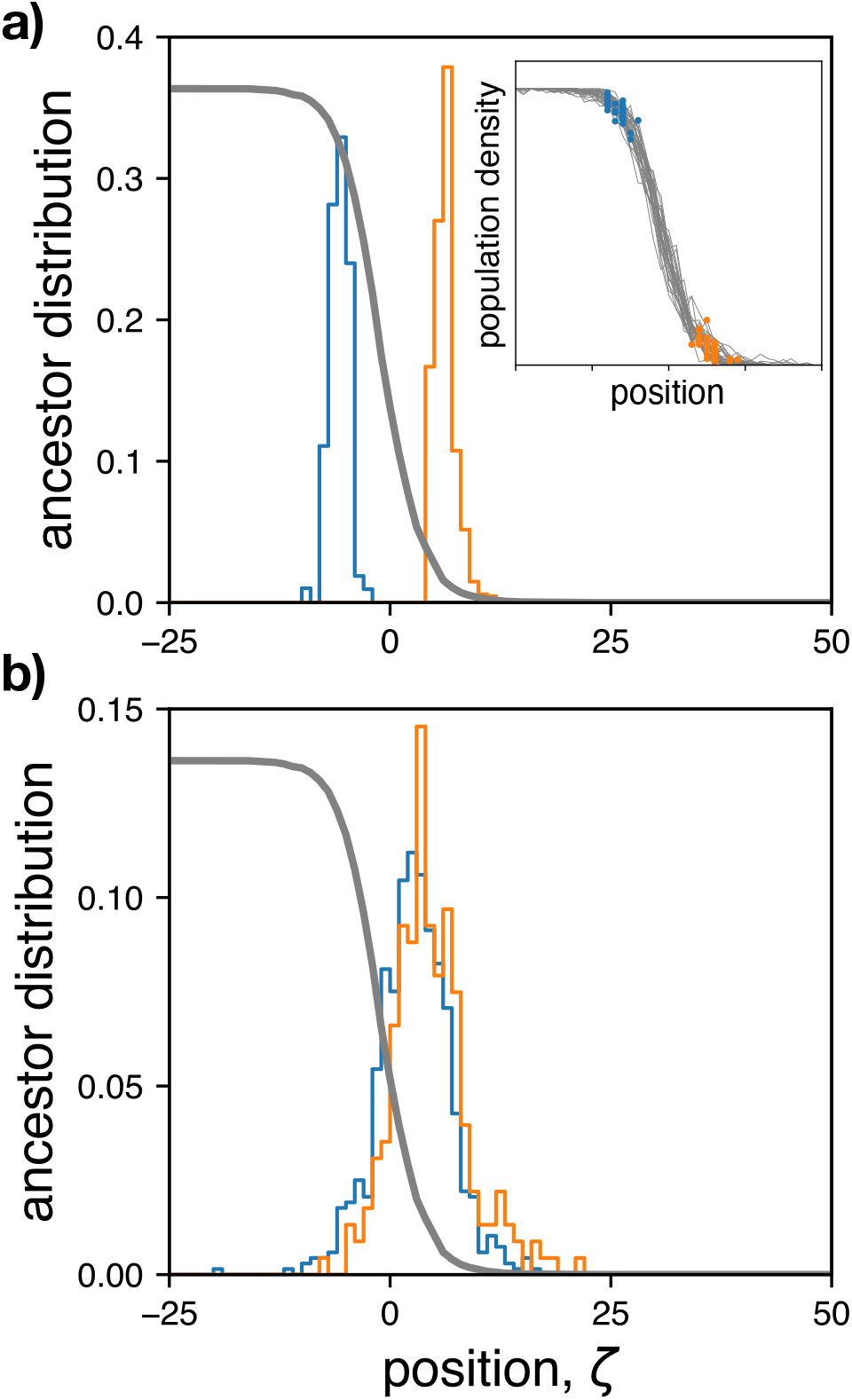
Genetic processes in spatially expanding populations are effectively well mixed on time scales larger than the mixing time of the front. **(a)** The distribution of initial locations from which the ancestor position was tracked backward in time. Due to the stochastic nature of the front the distribution of sampling locations has a finite width with respect to the average front profile shown in gray. (Inset) Shows final front from 30 independent runs used to generate histograms in the main panel. For each run, two subpopulations were chosen, one close to the bulk (blue) and another close to the edge of the front (orange), and the locations of their ancestors were recorded at different times in the past (see SI, Sec. V for exact sampling procedure). **(b)** Distribution of locations of ancestors of individuals shown in panel (a) from 100 generations in the past.

### Structure of genealogies

We performed simulations using three levels of cooperativity that are expected to lead to qualitative differences in the genealogies because they correspond to pulled, semi-pushed and fully pushed expansions. The genealogy of the population was obtained using the procedure described in the Methods section. The examples of these genealogies shown in Fig. 3 have the qualitative features predicted by the theory. In fully pushed expansions genealogies have only pairwise mergers, whereas semi-pushed and pulled expansions show several examples of multiple mergers. Moreover, the genealogies in pulled expansions appear highly skewed, with most mergers occurring on one side of the tree, while in fully-pushed expansions branching is more symmetric. These features are consistent with our hypothesis that cooperativity drives the transition from the Bolthausen-Sznitman to the Kingman coalescent.

**Figure 3:**
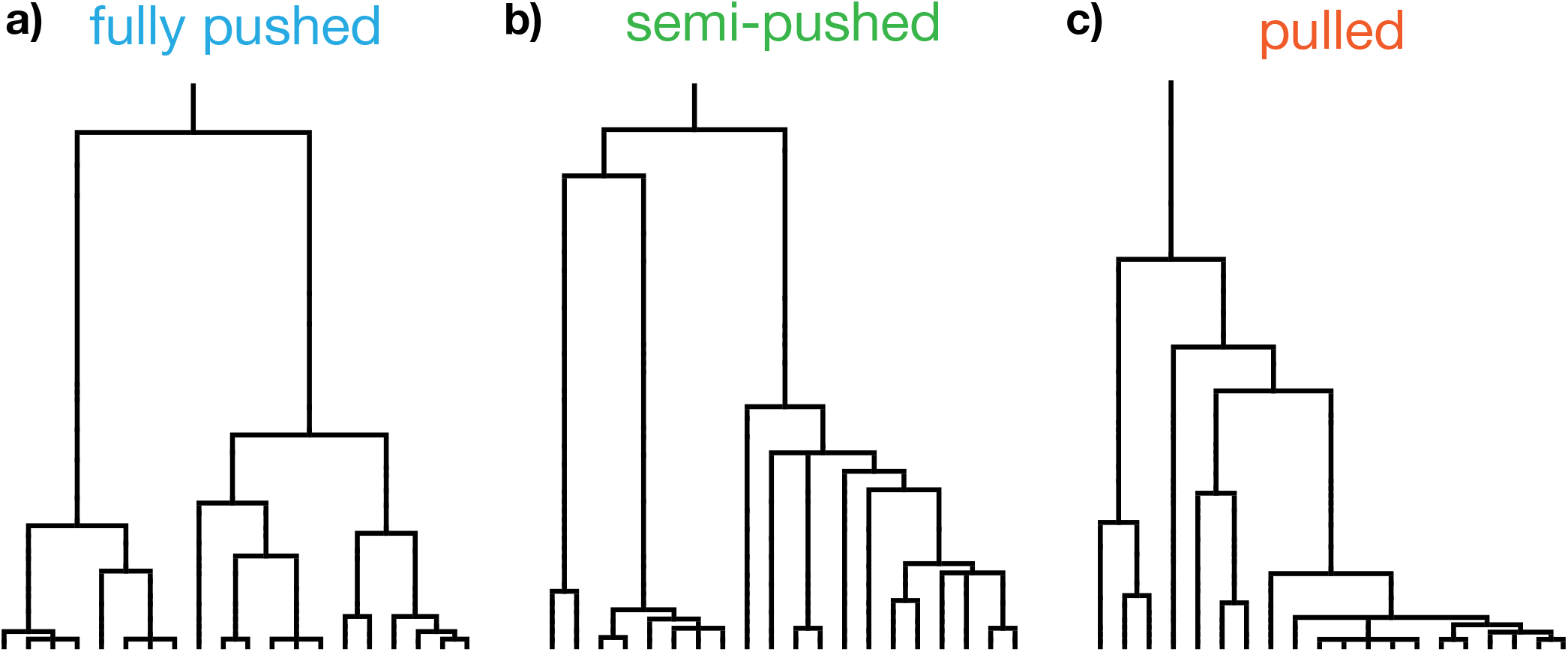
The genealogical tree of spatially expanding populations changes as the expansion transitions from pulled to pushed. Sample genealogies from fully pushed **(a)**, semi-pushed **(b)**, and pulled **(c)** expansions are shown. These trees were generated by randomly sampling 20 individuals from the first 15 occupied demes from the front, after fixation. For illustration purposes, we chose representative trees from our simulations that provided good visual clarity.

To get a more quantitative measure of the changes in topology of the genealogies during expansion, we calculated two summary statistics^3^ that can distinguish between coalescents: the site frequency spectrum (SFS), and the two-site frequency spectrum (2-SFS) [54,55]. We found that both SFS and 2-SFS supported our hypothesis that genealogies change from the Kingman to a non-Kingman coalescent at the transition between fully pushed and semi-pushed expansions. Because it is simpler to quantitatively test the SFS against the theoretical predictions, we report these results in the main text and refer the interested reader to Sec. III of the SI for the analysis of the 2-SFS.

The SFS provides a histogram of the number of sites in the genome that have a given frequency of mutations in the sample. Assuming mutation rates are constant throughout the genome, the SFS is the mean length of internal branches with a given number of terminal branches (leaves) [7,56]. We are particularly interested in the shape of SFS for high-frequency mutations (allele frequencies *f* ≈ 1) because SFS is qualitatively different between the Kingman and the Bolthausen-Sznitman coalescent in this regime.

High-frequency mutations occur on internal branches that have a large number of leaves. Genealogies with such mutations are highly skewed because one branch can contain the majority of leaves. Skewed trees are unlikely in the Kingman coalescent because each pairwise merger joins lineages randomly, independent of the number of their leaves. Thus SFS monotonically decays with the mutant frequency. In contrast, SFS for the Bolthausen-Sznitman coalescent is expected to have an uptick at high f because there is a high chance of nearly all lineages coalescing at a single multiple merger. Consistent with our hypothesis, we indeed find a monotonic SFS for fully pushed expansions (Fig. 4a), while semi-pushed and pulled expansions display the uptick at high allele counts characteristic of coalescents with multiple mergers (Fig. 4b, c). Moreover, both fully pushed and semi-pushed expansion SFS agree quantitatively with the predictions from the Kingman coalescent and the Beta-coalescent with *β* = 1.5, respectively (see SI, Sec. III for details). In the case of pulled expansions, we find the quantitative agreement is less good, which we believe is due to the very long relaxation times required to reach steady-state in the pulled regime (see SI, Sec. II). Nevertheless, taken together, these results clearly establish that the genealogies of the three expansion classes have distinct topologies.

**Figure 4:**
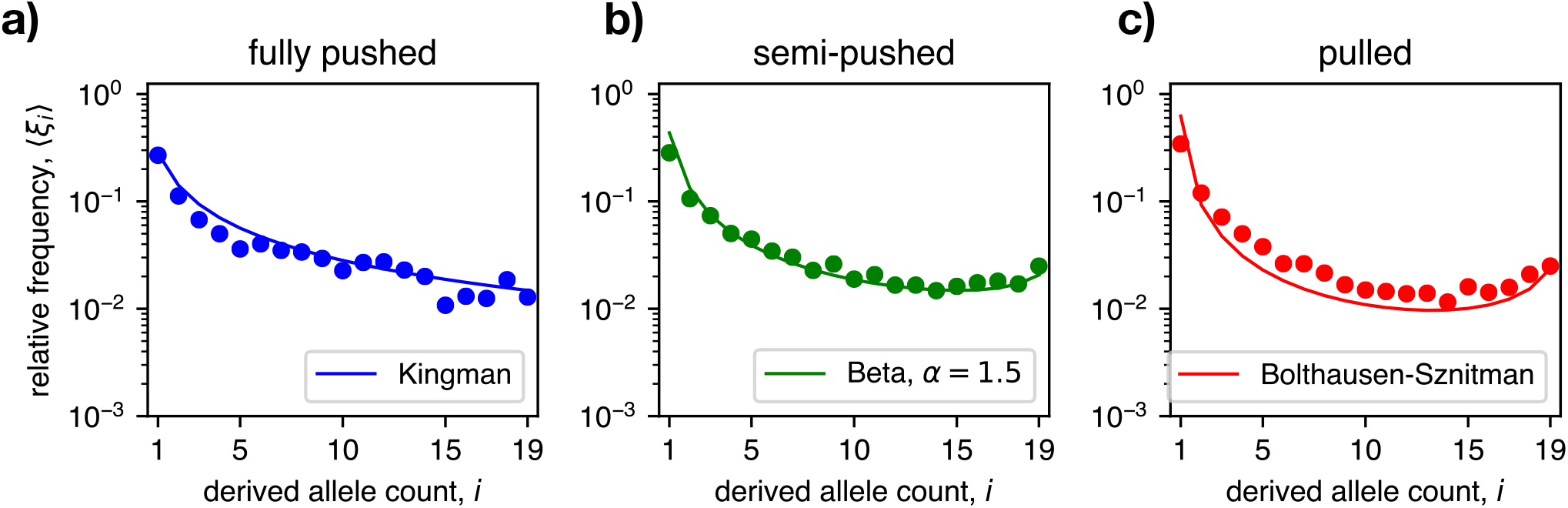
The site frequency spectrum of genealogies reveals differences between pulled and pushed waves. Approximately 100 trees were recorded from simulations of fully pushed **(a)**, semi-pushed **(b)**, and pulled **(c)** expansions, respectively. Each full genealogy was sampled 10 times using a sample size of 20 individuals chosen from the front (see SI, Sec V for sampling procedure). The resulting SFS, averaged over samples and simulations, is shown with colored dots. The solid line shows the exact predictions for the SFS in each regime (see SI, Sec. V for details).

## Theoretical results

### Descendant distribution in stochastic fronts

To develop an intuitive understanding of how genealogies emerge in range expansions, we developed a theoretical framework based on continuous reaction-diffusion equations. In this framework, it is easier to examine the dynamics of clones forward in time and relate the expansion of these clones to mergers in the genealogy. Previous work has shown that the frequency of a subpopulation *f_i_*(*t,ζ*) within the front changes according to the following equation [44,51]:

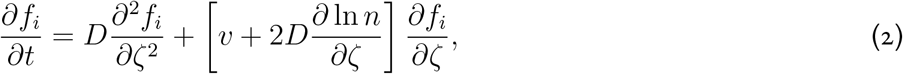

where *D* is the effective diffusion constant which describes the migration of individuals, *v* is the velocity of the front, *n*(*ζ*) is the population density, and *ζ = x* — *vt* is the position along the front in the comoving reference frame.

From (2) we can calculate the distribution of descendants from a single individual at some position *ζ*_0_ as *t* → ∞. In Sec. I of the SI, we show that this distribution has a time-independent form *f_i_*(*t, ζ*) ≈ *u*(*ζ*_0_) after sometime *O*(*τ_m_*), which we denote as the “mixing time” of the front. As a result, on time scales longer than *τ_m_* the distribution of surviving clones *f_i_*(*t,ζ*) loses all spatial information and *u* is simply proportional to the reproductive success of the ancestor.

Because *u* greatly varies with *ζ*_0_, individuals at different locations can have wildly different reproductive values *W*, which are determined by their average number of offspring after the mixing time [57,58]. We can invert this dependence and consider *ζ* (*W*)—i.e., find the location of the initial individual with a given reproductive value. It is then straightforward to compute the probability distribution for *W* by finding the number of organisms present at *ζ*(*W*). Mathematically, this is accomplished by the following change of variables: 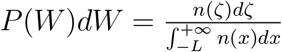. In Sec. I of the SI, we use this change of variables to calculate *P (W*) explicitly and find that it has a power law tail of the form

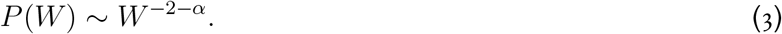

### The origin of different topologies

The exponent *α* is calculated exactly and depends only on *v/v_F_*, the ratio between the actual expansion velocity and the velocity that would occur in the absence of positive feedback 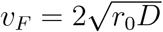:

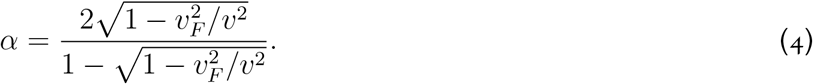

Note that the specific form of the density dependence in the growth and dispersal rates does not enter (4). In fact, all of our analyses have been carried out for an arbitrary model with short-range dispersal. Thus, the tails of *P*(*W*) are universal and depend on a single, easy-to-measure parameter [59].

For high cooperativity, when *v/v_F_* is greater than a critical value 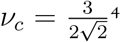, the exponent *α* is greater than one and the variance of *W* is finite. Therefore, the clone frequencies only change by small amounts each generation and genealogies are described by the Kingman coalescent [60]. For intermediate values of cooperativity, defined by 1 < *v/v_F_* < *ν_c_*, the exponent *α* is less than one and the variance of *P*(*W*) diverges. This leads to occasional large jumps in clone frequencies and the appearance of multiple mergers in the coalescent [47]. Finally, when *v/v_F_* = 1, we have *α* = 0 and *P*(*W*) ~ *W*^-2^, which leads to a Bolthausen-Sznitman coalescent when the process is viewed backward in time [47, 61].

To verify the change in descendant distribution predicted by theory, we measured clone sizes during range expansions in simulations. Direct measurements of *P*(*W*) are challenging because the distribution emerges only over a time scale of *O*(*τ_m_*), which we cannot determine precisely. However, we can circumvent this problem in two limits: on short time scales, on the order of a few multiples of *τ_m_*, and on long time scales, when the population comprises two clones. In the first limit, we can consider all individuals at the front at some initial time as clones of size one. As the front expands, some clones go extinct while others increase in size. For short time scales (comparable to *τ_m_*), clone sizes are small and each can be modeled as independent branching processes. In the second limit, we can track the dynamics of a population with only two clones—which we can think of as two alleles. As both alleles are neutral, the dynamics can be described by the frequency of one of them, which changes according to a Fleming-Viot process [61, 62].

The branching process calculation makes two testable predictions about the clone size distributions. First, the average size of a surviving clone 〈*W*〉_+_ increases as *t*^1/*α*^. Second, the probability to observe a clone *s* times larger than the average clone decays as *e*^-*s*^ for *P*(*W*) with a finite variance and as *s*^-1-*α*^ when *α* < 1. In the SI we show the results of simulations for fully pushed expansions agree well with these predictions (Fig. S4). Outside of the fully pushed regime, we see a broadening in the clone size distribution which is inconsistent with the exponential prediction for a short-tailed descendant distribution (Fig. S4). However, due to the large carrying capacities required to allow for the relaxation of the transient dynamics in the semi-pushed and pulled regimes, we were not able to quantitatively verify the expected power law for *F*(*s*).

The simulations of the Fleming-Viot process were more efficient and allowed us to demonstrate a quantitative agreement with our theoretical predictions. Specifically, we started forward-in-time simulations with two clones of equal abundance and monitored the frequency *f* of one of the clones. Conditioned on having both clones present, the probability *P* (*f*) of observing a particular clone frequency approaches a steady state in simulations and can also be computed analytically. For the Kingman coalescent, *P*(*f*) = 1 [63] while for the Bolthausen-Sznitman coalescent 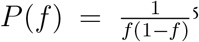. Our simulations mach both of these predictions (Fig. 5a-c).

**Figure 5:**
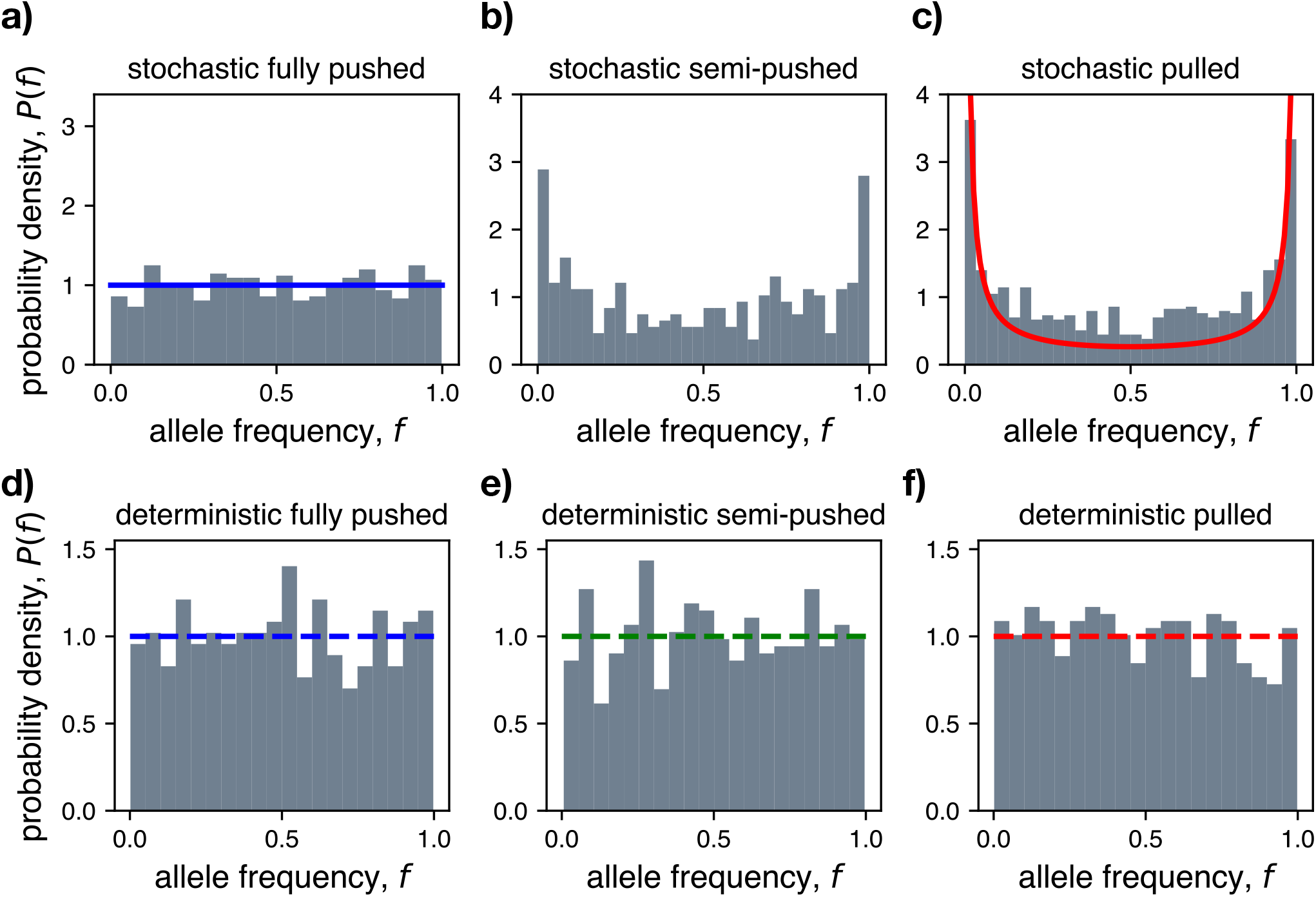
Deterministic front approximation fails to capture full range of coalescent topologies. **(a-c)** Shows long-time distribution of the frequency of one allele in stochastic two-allele simulations for each expansion type. Histogram of allele frequencies are shown in gray and the theoretical predictions assuming the Kingman (panel a, blue) and Bolthausen-Sznitman (panel c, red) coalescent are show with solid lines. **(d-f)** Same as above, but for simulations with a deterministic front. The dashed lines show the theoretical predictions, which are now given by the Kingman coalescent. For each panel we ran 10^3^ simulations and report the distribution of allele frequencies at a fixed time after the distribution becomes quasi-stationary.

### The role of fluctuations in population density at the front

All of our results so far explicitly account for demographic fluctuations at the front. However, most studies of range expansions have ignored demographic fluctuations, either because of the mathematical difficulties they introduce or because their effects were thought to be small [32, 64, 65]. To understand to what extent density fluctuations influence the dynamics at the front, we performed simulations in which the total population density was updated deterministically, while still allowing for genetic drift by stochastically sampling the front composition. In all simulations, we found that genealogies matched the Kingman coalescent (Fig. 5d-f).

This unexpected result can be explained by considering the effect of deterministic population dynamics on the descendant distribution at the front. In the Methods section, we show that the deterministic approximation leads to a finite variance in *P*(*W*), through a cutoff *W_c_* corresponding to the maximum reproductive value at the front. We also show that the cutoff *W_c_* scales sublinearly with the carrying capacity *N* (see SI, Sec. I). This implies that the fraction of lineages which can merge in one event in the limit of large *N* goes to zero. As large merger events are suppressed, we expect all genealogies to converge to the Kingman coalescent. Thus, demographic fluctuations play a crucial role in the emergence of non-Kingman coalescents at the front.

## Discussion

Many species, from microbes [66, 67] to humans [23], have undergone expansions in their history and many are currently expanding due to globalization [68, 69] and climate change [27, 70]. Previous work has demonstrated that range expansions reduce the amount of genetic diversity in the population [32, 64, 71, 72] and allows for some alleles to become dominant, through a process known as gene surfing [51, 65, 73]. However, underneath the overall decrease in diversity many patterns can be found which are still not well understood.

Evolutionary dynamics during range expansions vary greatly depending on how much demographic fluctuations and genetic drift at the leading edge influence future generations [44]. This dependence is captured by a single parameter *v/v_F_*. This ratio of the actual expansion velocity to the velocity that would occur without density dependence quantifies the degree of cooperativity (or positive feedback) in growth and dispersal. When this parameter is large, the front makes a small contribution to the rate of expansion and allele frequencies change slowly. When *v/v_F_* is close to one, expansion proceeds primarily via a highly stochastic advancement of the population edge.

We showed that these differences in evolutionary dynamics are captured by a simple and intuitive model, which describes the front as an effective well-mixed population with broad distribution of reproductive values. As *v/v_F_* decreases, the descendant distribution becomes broader until, at a critical value, the variance diverges—this signals the transition from the Kingman to a non-Kingman coalescent. As *v/v_F_* decreases further, the distribution broadens until a Bolthausen-Sznitman coalescent is reached.

Density fluctuations are essential for all of our results. When these fluctuations were ignored, all genealogies were described by the Kingman coalescent, as predicted from the serial bottleneck view [32]. More sophisticated models have attempted to replace the effects of demographic fluctuations by a cutoff at *n(ζ_c_) =* 1. While such a cutoff is appropriate for pulled expansions, for others it is not [74]. It has recently been discovered that for semi-pushed and fully pushed expansions, a different cutoff, which depends on *v/v_F_*, should be used [44]. There, *quantitative* changes in the rate of diversity loss were found when the wrong cutoff was used. Here, we found the choice of cutoff leads to *qualitative* changes in the genealogies. Thus, any theory that hopes to predict the dynamics of expansions needs to account for fluctuations in the position and shape of the front.

Our results provide a universal framework to link genetic diversity at the front to ecological dynamics. This framework can be used to infer the importance of density feedback in growth and dispersal or to predict evolution during range expansions. More importantly, the mechanism presented here provides a generic explanation for the skewed genealogies commonly observed in empirical studies [19, 75–77]. Previously such genealogies were attributed to either very strong selection or sweepstakes reproduction [19, 77], both of which could be less common than range expansions. Nevertheless, natural populations are both spatially structured and under various selection pressures—integrating both aspects is required for developing a complete theory of genealogical trees.

## Acknowledgments

The authors would like to thank Benjamin H. Good for helpful discussions and comments. K.S.K. and G.B. were partially supported by the Simons Foundation Grant #409704; K.S.K. also acknowledges the support by the Research Corporation for Science Advancement through Cottrell Scholar Award #24010 and by NIGMS through grant #1R01GM138530-01. G.B. was also partly supported by the Simons Foundation Postdoctoral Fellowship Award #730295. O.H. acknowledges the support from the Simons Foundation Grant #409704, the National Science Foundation Career Award #1555330, and the National Institute of General Medical Sciences of the National Institutes of Health Award R01GM115851. Simulations were carried out on the Boston University Shared Computing Cluster.

## Supplemental Information

### I. Forward in time dynamics

In this section, we show how the genealogy of an expanding population can be mapped to an effective well-mixed population with a broad descendant distribution. We only consider the case of density-independent migration here, but the argument is analogous when *D* depends on *n*. We consider a population with density *n*(*t, x*) described by

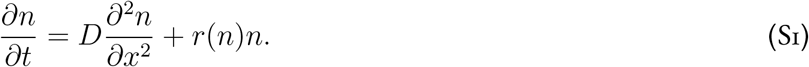

We assume the population is comprised of *m* neutral subtypes with relative fractions *f_i_*(*t, x*) and 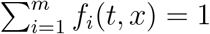. In the deterministic limit, it is then easy to show that *f_i_*(*t, x*) obey the following equation [1, 2]: descendant

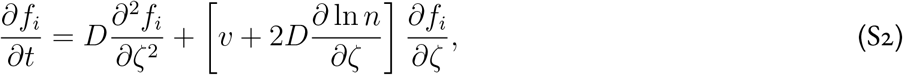

where *ζ* ≡ *x – vt* is the spatial coordinate in the comoving reference frame. We can write the general solution for *f* (*t, ζ*) as:

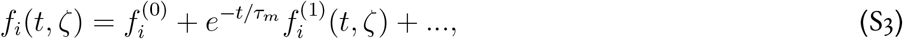

where we have kept the two eigenvectors of the operator in Eq. (S2) with the slowest decay times. For large *N*, the timescale *τ_m_* is smaller than the mean coalescence time *T_c_*, and represents the time for an arbitrary distribution of neutral alleles to “mix” with the other individuals at the front and reach its steady state distribution [3]. We will, therefore, refer to *τ_m_* as the mixing time of the front.

Previous work has shown that lim_t→∞_ *f* (*t, ζ*) = *u*(*ζ*), where *u*(*ζ*) is the fixation probability of a new mutant that originates at position *ζ* (see Sec. III of SI of Ref. [3] for an extensive discussion). The fixation probability can be calculated explicitly and has the following form:

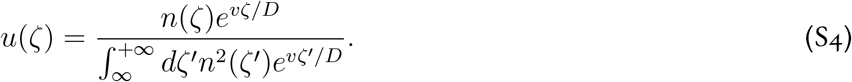

If we interpret *u*(*ζ*) as the fraction of descendants of an individual at position *ζ* in the whole population, and use *f* (*t,ζ*) ≈ *f*^(0)^ for *t* ≳ *τ_m_*, we can think of the population of the wave as an effective well-mixed population with a broad descendant distribution *P* (*W*), and generation time *τ_m_*. The relation between the number of descendants W can then be computed using

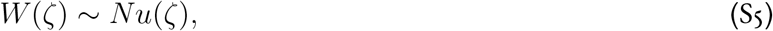

where carrying capacity *N* is a necessary conversion factor because *u*(*ζ*) is a probability, and strictly less than one.

We can compute the descendant distribution by using

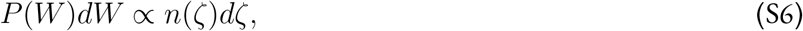

to eliminate the position ζ. The result then reads

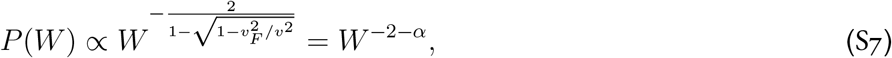

where

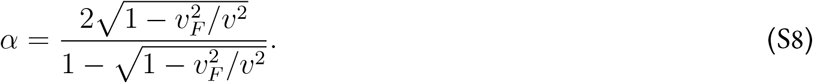

For pulled waves, *v/v_F_* = 1 andwehave *P* (*W*) ∝ *W*^-2^. This distribution has a divergent mean and leads to Bolthausen-Sznitman coalescent [4]. In the semi-pushed region, 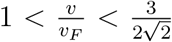 and the descendant distribution changes continuously from *W*^-2^ to *W*^-3^. Finally, in fully-pushed waves, it decreases at least as fast as *W*^-3^. In this case, the population is described by a Kingman coalescent [5].

#### Cutoff in descendant distribution

The above argument applies for deterministic dynamics of *n*(*t, x*). However, it is not clear whether models with stochastic migration and growth behave the same way. Previously, we argued that stochasticity can be incorporated by using an effective cutoff in the population density [3], which for semi-pushed waves is given by

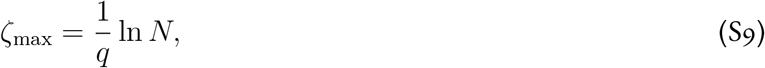

where 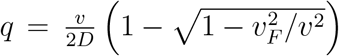. Using this cutoff we can compute the maximum fraction of descendants in the population:

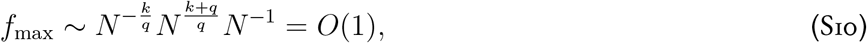

where we have defined 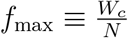. The above result shows that the cutoff does not depend on *N*, and does not influence the properties of coalescent for *u* ≪ 1. However, this does not exclude a finite cutoff at some *u_c_* < 1, which would change the frequency of very rare fluctuations, were a fraction ≲ 1 of lineages merge during one generation. A more detailed calculation is needed to check for this.

#### Deterministic waves

If we apply the same reasoning when *n*(*t, x*) is discrete but changes deterministically, we get a very different answer. In this case, we have a cutoff at *n* = 1, which occurs at

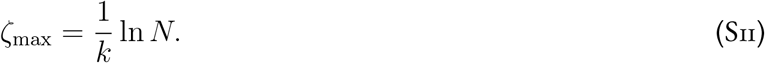

In the semi-pushed regime, this gives a maximum value for *W*:

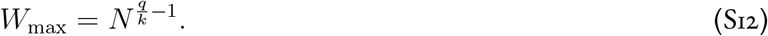

Looking backward in time, we can express the cutoff at *f*_max_ in terms of the largest fraction of lineages in the population that can coalesce into one individual over a timescale of *τ_m_*. The results reads

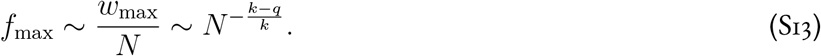

Looking backward in time, *u*_max_ represents the largest fraction of lineages in the population that can coalesce over the generation time *τ_m_*. Since for pushed waves *q* < *k*, this shows that in the limit of *N* → ∞

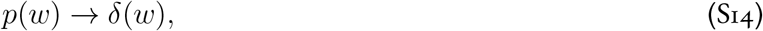

and the genealogical tree converges to a Kingman coalescent. This prediction is particularly striking since we have shown that *T_c_* still has a power law scaling with *N* even for deterministic fronts [3].

#### Distribution of allele fraction

We can better understand the coalescent structure during expansions by studying the distribution of allele frequencies f for at long times. For a Kingman coalescent, the allele frequency is given by the classic result of Kimura [6]:

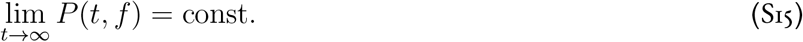

At the other extreme, the Bolthausen-Sznitman coalescent is the dual of a jump-advection process, with the distribution [4]

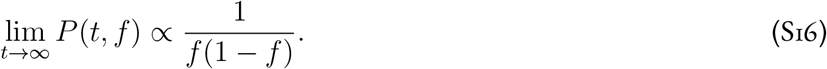

### II. Effective clone size distribution

In this section we calculate the clone size distribution at the front on times scales much longer than *τ_m_*, by approximating the process in the effective well-mixed population by a branching process. The probability distribution can be obtained analytically for 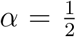 and when the number of descendants has a finite variance. We briefly review the history and the relevant references and then summarize the key results explaining briefly how they can be derived.

#### Relevant literature

Branching processes were first studied by Watson and Galton to describe the dynamics of British surnames [7]; therefore they are often referred to as Galton-Watson processes. Branching processes have been applied to a number of fields including branching of neutrons in nuclear reactions, population genetics, earthquakes, chemical reaction, birth-death processes, shot noise, and many others. The monograph by Harris [8] contains the historical details and detailed mathematical treatment of simple and generalized branching processes together with several applications. A simpler and more limited exposition can be found in Ref. [9]. A summary of the early progress in branching processes can be found in Ref. [10]. Branching processes were also called multiplicative processes possibly because of the application to the nuclear reactions; see [11].

The full solution for the branching process was developed by a great number of scientists who calculated different properties under different assumptions. Some of the key results that are relevant for us were obtained in Ref. [12, 13]. The approach taken in the latter reference is very close to how a physicist would approach this problem and our discussion closely follows that of Ref. [13]. More recently, branching processes have been used in the study of avalanches and total popularity on networks [14,15]. These references extend the classical results to compute the integral of the number of organisms over time for surviving lineages, i.e. avalanche size. On the mathematical sized, branching processes can be studied in the continuum limit, which is known as continuous state branching processes. This description is equivalent to a Levy process with a time change. All of the results, however, can be derived from the discrete number of individuals by taking the continuum limit [16–18].

#### Problem formulation and general solution

We consider a continuous time version of the branching process since it is simpler. The probability to observe *n* individuals at time *t* is denoted as *p_n_*(*t*). Unless specified otherwise, we assume that *p_n_*(0) = *δ*_*n*,1_. The probability to leave *k* descendants is *q_k_*.

The master equation reads

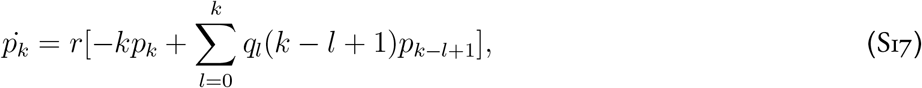

where *r* is the branching rate. Since *r* only enters the problem through the time scale, we set *r* = 1 in the following.

Note that the transition rates are proportional to the number of individuals since each can reproduce. The fact +1 in the last term accounts for the fact that the reproducing individual dies.

The master equation can be solved using generating functions. We denote the generating functions for *p_n_* and *q_k_* by *P* and *Q* respectively:

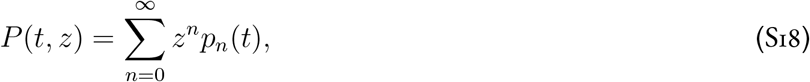

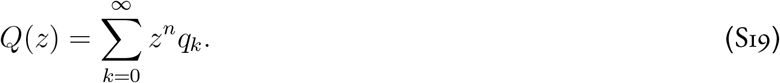

Upon differentiating Eq. (S18) with time and using Eq. (S17), we obtain

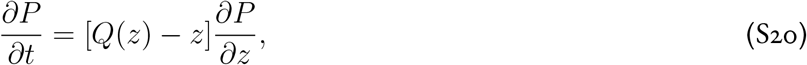

which can be solved using the method of characteristics. Assuming that we start with one individual, *P*(0, *z*) = *z*, and the implicit solution of Eq. (S20) reads

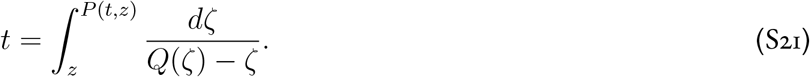

This equation serves as the basis of our analysis in the rest of this summary.

Before proceeding with the analysis, however, we point out that many references study branching processes from a different starting point. Consider how the population can change in a short time *dt* at the start of the process when there is only one individual (similar to backward Kolmogorov equation). With probability 1 — *dt*, nothing happens and the generating function remains unchanged. With probability dt the organism reproduces and leaves *k* descendants with probability *q_k_*. After that we also have a branching process that lasts time *t*, but starts with *k* individuals. Since individuals are independent the generating function for the sum of their progenies is the product of the generating functions for each starting organism. In other words, we obtain

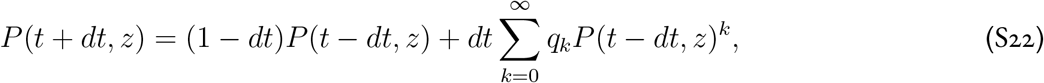

which simplifies to

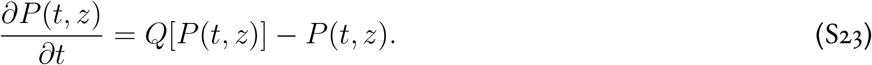

It is easy to see by direct substitution that the implicit solution from Eq. (S21) satisfies Eq. (S23). The direct analysis of Eq. (S23) and its discrete-time analog involves functional equations and recurrences, which are more cumbersome than the implicit solution obtained above.

#### Asymptotic analysis

When the integral in Eq. (S21) can be evaluated one can obtain *P*(*t, z*) directly. For a general *Q*(*z*), we focus on long time limit. In this limit, *t* → +∞ and the integral must diverge. Therefore, the long time behavior of *P*(*t, z*) is controlled by the root *z_c_* of *Q*(*z_c_*) = *z_c_* and the behavior of *Q*(*z*) around *z_c_*.

It is easy to show that *z_c_* > 1 when the mean number of descendants 〈*k*〉 = *Q*^!^(1) < 1. In this case, *P*(*t,z*) approaches 1 exponentially fast, which corresponds to guaranteed extinction. Note that any generating function needs to be less or equal to one for |*z*| ≤ 1.

When 〈*k*〉 = *Q*’(1) > 1, *z_c_* < 1. In this case, the process has a finite probability to survive, which is given by 1 — *z_c_*. The population size of surviving realizations grows exponentially with time at a rate given by 〈*k*〉 — 1. More refined results can be obtained by expanding *Q*(*z*) in Taylor series around *z_c_*.

When 〈*k*〉 = *Q*’(1) = 1, we have a critical branching process. This is the case that we will focus on in the following. In this case *z_c_* =1 and the behavior of *P*(*t, z*) depends on the behavior of *Q*(*z*) around *z* = 1. If 〈*k*^2^〉 exists, *Q*(*z*) has a second derivative at *z* =1 and can be approximated by *Q*(*z*) = *z* + 1/2*Q*”(1)(1 – *z*)^2^. If the variance is infinite, then *Q*(*z*) is not analytic around *z* =1. We argue below that, when the number of descendants is distributed according to a power law, *Q*(*z*) = *z* + *g*(1 — *z*)^1+*α*^ with *α* ∈ (0,1].

In the next two sections, we evaluate the integral in Eq. (S21) using the approximations for *Q*(*z*) to obtain the long time asymptotics of *P*(*t, z*).

#### Critical branching process with finite variance

Upon substituting 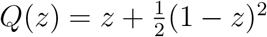 into Eq. (S21) and evaluating the integral, we obtain

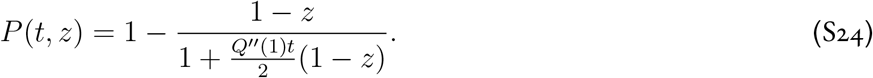

The survival probability is given by

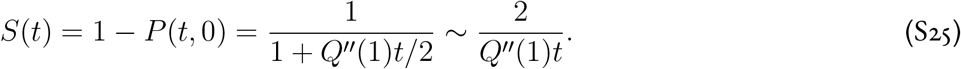

The average size of a surviving lineage 〈*n*〉_+_(*t*) should be such that 〈*n*(*t*)〉 = 1. Therefore

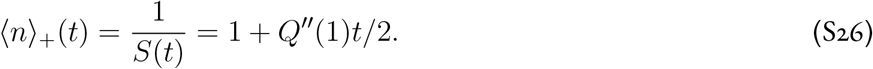

To obtain *p_n_*(*t*), we expand *P*(*t, z*) in Taylor series around *z* = 0. The result for *n* > 0 reads

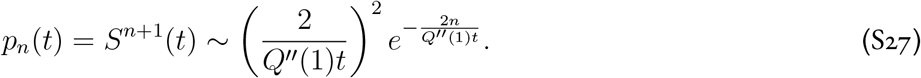

The above expression can be recast in a simpler form by normalizing the population size by the expected population size of surviving realizations. Specifically, we let *y* = *n*/〈*n*〉_+_, which also affects the normalization constant, and divide *p_n_* by *S* (*t*) since we consider only surviving realizations. The distribution of scaled population sizes is then described by the following probability density function:

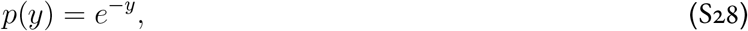

where we omitted the time since the equation corresponds to the limit *t* → +∞. This relationship can also be derived in a more formal and general way that we describe below.

#### Continuum limit from generating function

Given *P*(*t, z*) how can we obtain *p*(*t, y*)? First, notice that

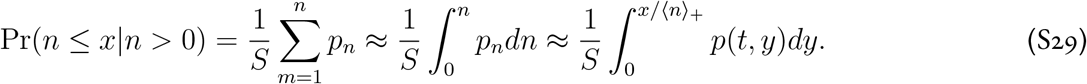

Therefore

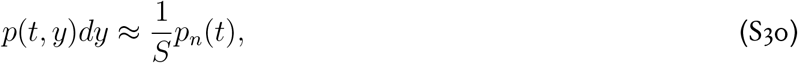

and

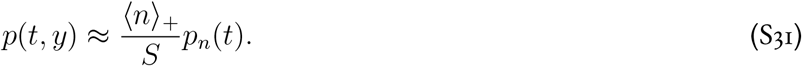

Then, we can relate the generating function *P*(*t, z*) to the moment generating function of *p*(*t, y*):

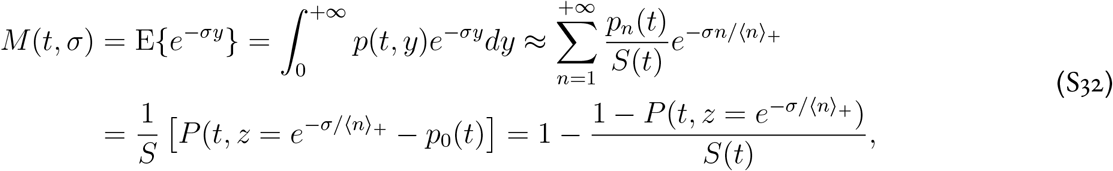

where we used *p*_0_ = 1 — *S*. One can then obtain *p*(*t,y*) via an inverse Laplace transform of *M*(*t,σ*). Note that for the critical branching process 〈*n*〉_+_ = 1/*S*.

Since it is convenient to summarize simulation results in terms of the complementary (reverse) cumulative distribution *c*(*t, y*), we also derive the connection between *P*(*t, z*) and the Laplace transform of *c*(*t, y*):

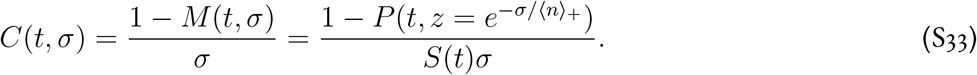

We can apply this result to the branching process with finite variance to obtain the long time limit of *c*(*t, y*) as follows:

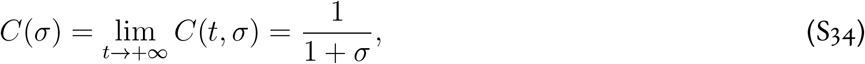

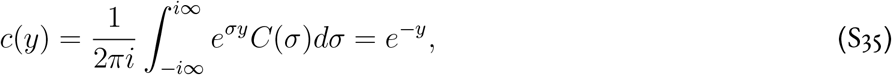

which indeed describes the complementary cumulative distribution for *p*(*y*).

#### Power-law tails and the behavior of the generating function

Before repeating the analysis above for distributions of the number of descendants *q_k_* with diverging variance, we briefly discuss the connection between the power law tail of *q_k_* and the singularity of *Q*(*z*) at *z* = 1. As a reminder, we focus only on critical branching processes with 〈*k*〉 = 1 and only on *q_k_ ~ k*^-2-*α*^ for large *k*. Under these assumptions,

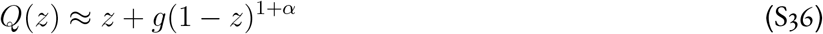

around *z* = 1.

To see this, one can compute *q_k_* from the equations above by expanding *Q*(*z*) in Taylor series around *z* = 0 using the Cauchy formula for the derivatives:

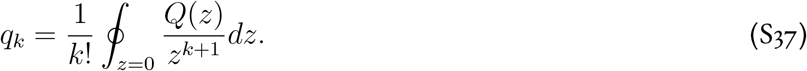

This integral can be evaluated by taking the branch cut along (1, +∞), moving the contour to hug the branch cut, changing the integration variable from *x* to *e^p^*, and observing that only *p* ≲ 1/*k*. The final result reads

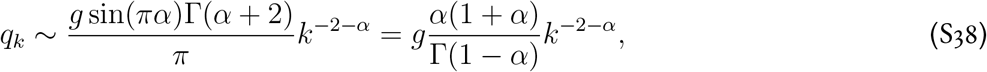

where Γ(*x*) is the Gamma function.

Another way to derive the relationship is to choose a specific form of *q_k_*. A convenient choice is *q_k_ = k*^-(2+*α*)^/*ζ* (1 + *α*) for *k* > 0 and *q*_0_ = 1 — *ζ*(2 + *α*)/*ζ*(1 + *α*), where *ζ*(·) is the Riemann zeta function. Note that this choice satisfies both the normalization condition and the requirement that the average number of descendants equals to one. It is easy to show via a Taylor expansion around *z* = 0 that the corresponding generating function is given by

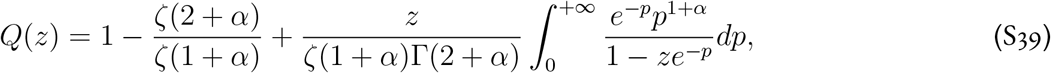

where the last term without the zeta function is known as Li_2+*α*_ (·), polylogarithm of order 2 + *α*. The asymptotics of *Q*(*z*) can be directly extracted from this integral representation or from the asymptotics of the polygarithm.

#### Critical branching process with diverging variance

To find *P*(*t, z*), we substitute the approximation for *Q*(*z*) (Eq. (S36)) into the implicit solution given by Eq. (S21). The result reads

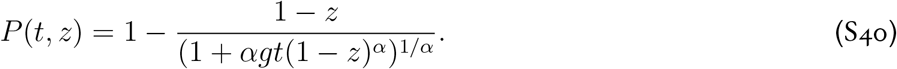

This expression contains all the information that we need. In particular, one can pass to a continuum limit and obtain *C*(*t,σ*) and *c*(*t, y*). Inverse Laplace transform can be evaluated by moving the integration contour to hug the branch cut (—∞, 0). Below, we consider a few special cases where the calculations are particularly simple and provide additional insight.

The survival probability and the average size of the surviving population are given by

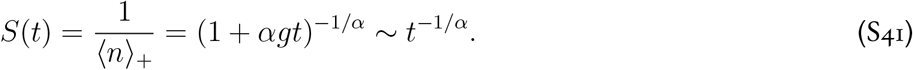

Note that the relevant time scale is 1/(*αg*), which becomes *ζ*(1 + *α*)(1 + *α*)/Γ(1 – *α*). The latter expression scales as 1/*α* for *α* → 0. Thus, one should expect very long transient dynamics for small *α*.

The long time limit for *C*(*t, σ*) is given by

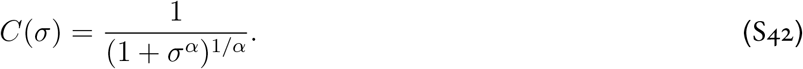

The inverse Laplace transform yields the following asymptotics

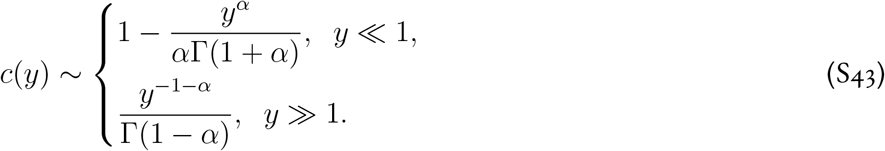

The asymptotics for *p*(*y*) are obtained by differentiation with respect to *y*.

For the special case of *α* = 1/2, one can obtain an analytic expression for *c*(*y*):

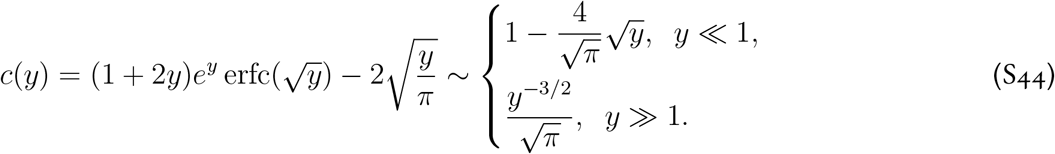

The small *y* asymptotics can also be derived directy from the generating function by expanding it in Taylor series around *z* = 0. This yields

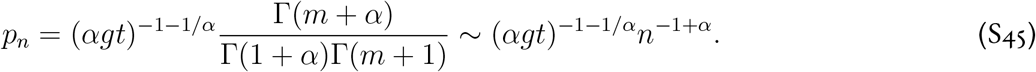

#### Clone sizes in a stationary process

Note that the results above might seem surprising at first. Most of the time branching processes are not conditioned on starting at a particular time. Instead, one assumes that the process restarts once extinction occurs. The sampling from such a stationary process gives more weight to processes that survived for a long time and therefore had proportionally large chance to be sampled.

It is easy to show that the distribution of the age *p*(*a*) of a process sampled at a random time is given by *S* (*t = a*). Indeed,

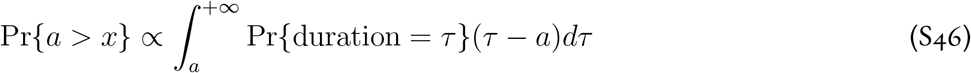

then

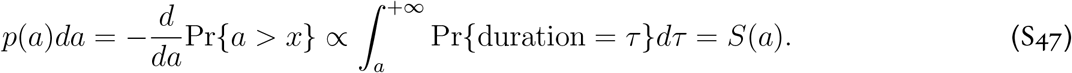

The probability to observe a population of size *n* is then given by

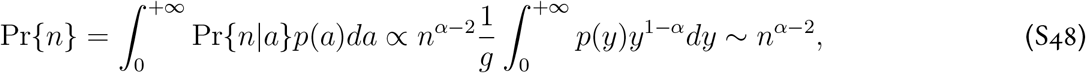

where we expressed Pr{*n|a*} as *p*(*y* = *n*/〈*n*〉_+_), i.e. using the probability distribution for the scaled population size defined in previous sections.

Equation (S48) agrees with the classical results for the neutral model for *α* = 1 and describes the tail of the site-frequency spectrum for the Kingman and Bolthausen-Sznitman coalescents.

### III. Summary statistics of ancestral trees

In this section we present other summary statistics we used to infer the topology of the ancestral trees obtained from simulations.

#### Theoretical background

Our analysis of the genealogies is based on the coalescent theory. The coalescent provides a model for the backwardin-time dynamics of lineages in a population without any internal structure^6^. Generally, such a model is completely described by the rates λ_*b,k*_ at which *k* out of *b* lineages merge. An important result shows that λ_*b,k*_ for any coalescent^7^ can be written in the following form:

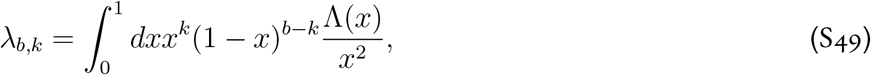

where 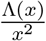 is the distribution of the merger sizes [5]. A few special choices of Λ(*x*) are worth noting. First, Λ(*x*) = *δ*(*x*) gives λ_*b*,2_ = 1 and λ_*b,k*_ = 0 for *k* > 2. This is the standard Kingman coalescent, where only pairwise mergers are allowed and their rate is constant. Another important model is the Bolthausen-Sznitman coalescent and is given by Λ(*x*) = 1. The merger rates in this case are 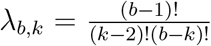, which implies that *k* = 2 and *k* = *b* mergers are equally likely and the most likely merger size is close to 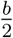. Such large merger events have been used to describe genealogies of populations under strong selection [24, 25]. Finally, one can interpolate between the two by using

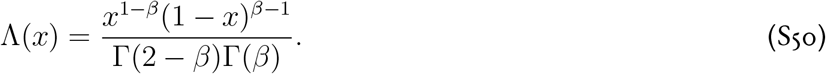

From (S50), it is easy to show that *β* =1 gives the Bolthausen-Sznitman coalescent and *β* = 2 gives the Kingman coalescent. The coalescent described by (S50) is known as the Beta-coalescent and many of its properties have been studied previously [5]. For our purposes, it is important to note that the Beta-coalescent describes the genealogies of highly fecund populations, in which the descendant-number distribution *P*(*W*) has a power law tail *P*(*W*) ~ *W*^-(1+*β*)^. We demonstrate in the Results section that the descendant-number distribution has a power law tail in range expansions, when viewed over a few generations. We make use of this fact to argue that genealogies in range expansions can generically be described by a Beta-coalescent.

The choice of Λ(*x*) in (S49) has important effects on the different summary statistics which we use to characterize genealogies. One such such statistic is the average time for two lineages to coalesce *T_c_*, whose scaling with the population size is highly dependent on the coalescent [5]. Thus, in the Kingman coalescent, *T_c_* is typically proportional to the population size, while in the Bolthausen-Sznitman coalescent it has much weaker logarithmic dependence. These distinct scaling regimes match our previous results, showing that *T_c_ ~ N* in fully pushed expansions, and *T_c_* ~ ln^3^ *N* in pulled expansions, with semi-pushed expansion having a sublinear power law dependence with a tunable exponent and lying in between [3]. Based on these results, we expect the genealogies in range expansions to be described by a continuum of Beta-coalescents as in (S50), spanning the range from the Kingman to the Bolthausen-Sznitman coalescent.

We also consider the site frequency spectrum (SFS), which corresponds to the set of lengths of branches *ξ_k_* subtending *ξ_k_* leaves, for all values of *k* ∈ [1, *n* — 1]. As with the total tree length, the exact SFS can be obtained recursively for small *n* [26]. Asymptotic results for large *n* are also known for (S50), but they converge slowly with sample size [26] and we will not use them here. Similar results can be obtained for 2-SFS, which represents the covariances between branch lengths [26].

#### Two-site frequency spectrum

While the SFS of the Beta- and Kingman coalescents are quite different as we have shown, relaxing the assumption of constant population size in the Kingman coalescent can lead to the site frequency spectra becoming more similar. Recently, it has been proposed that the two-site frequency spectrum (2-SFS) is a more robust measure to distinguish between Kingman and non-Kingman coalescents [27]. The 2-SFS, *p_n_*(*k, l*), for a sample size *n* is the number of pairs of sites which have derived allele counts *k* and l. For constant mutation rates, the 2-SFS can be derived from the genealogical tree—in this case *p_n_*(*k, l*) is proportional to the second moment of the length of branchs that subtend *k* and *l* leaves. In the case of the Kingman coalescent, the long branches near the common ancestor lead to a large number of sites which co-occur or split the tree in half. This explains the high values of the 2-SFS seen on the diagonals. In addition, pairwise branching of ancestral lineages constrain the topology further down tree, leading to anticorrelations between rare alleles (Fig. S1a). In contrast, coalescents with multiple mergers have shorter branches near the common ancestor, decreasing the density along the diagonal of the 2-SFS. The tree topology of coalescents with multiple mergers is also less constrained by early mergers, resulting in less pronounced negative correlations between rare alleles in the Beta-coalescent, and positive correlations in the Bolthausen-Sznitman coalescent (Fig. S1b, c).

We used the trees generated from simulations of fully pushed, semi-pushed, and pulled expansions to test 2-SFS against the theoretical predictions. We found that the patterns in the 2-SFS matched the theoretical predictions for Kingman, Beta-, and Bolthausen-Sznitman coalescents quite well (Fig. S1). In particular, fully pushed waves showed negative correlations outside of the main diagonals as expected, with correlations below the main diagonal smaller in absolute value than those above the main diagonal. Semi-pushed and pulled expansions, on the other hand, showed signatures of multiple mergers in the form of an increase of correlations below the main diagonal, and higher positive correlations on the off-diagonal, especially in the case of pulled expansions.

**Figure S1:**
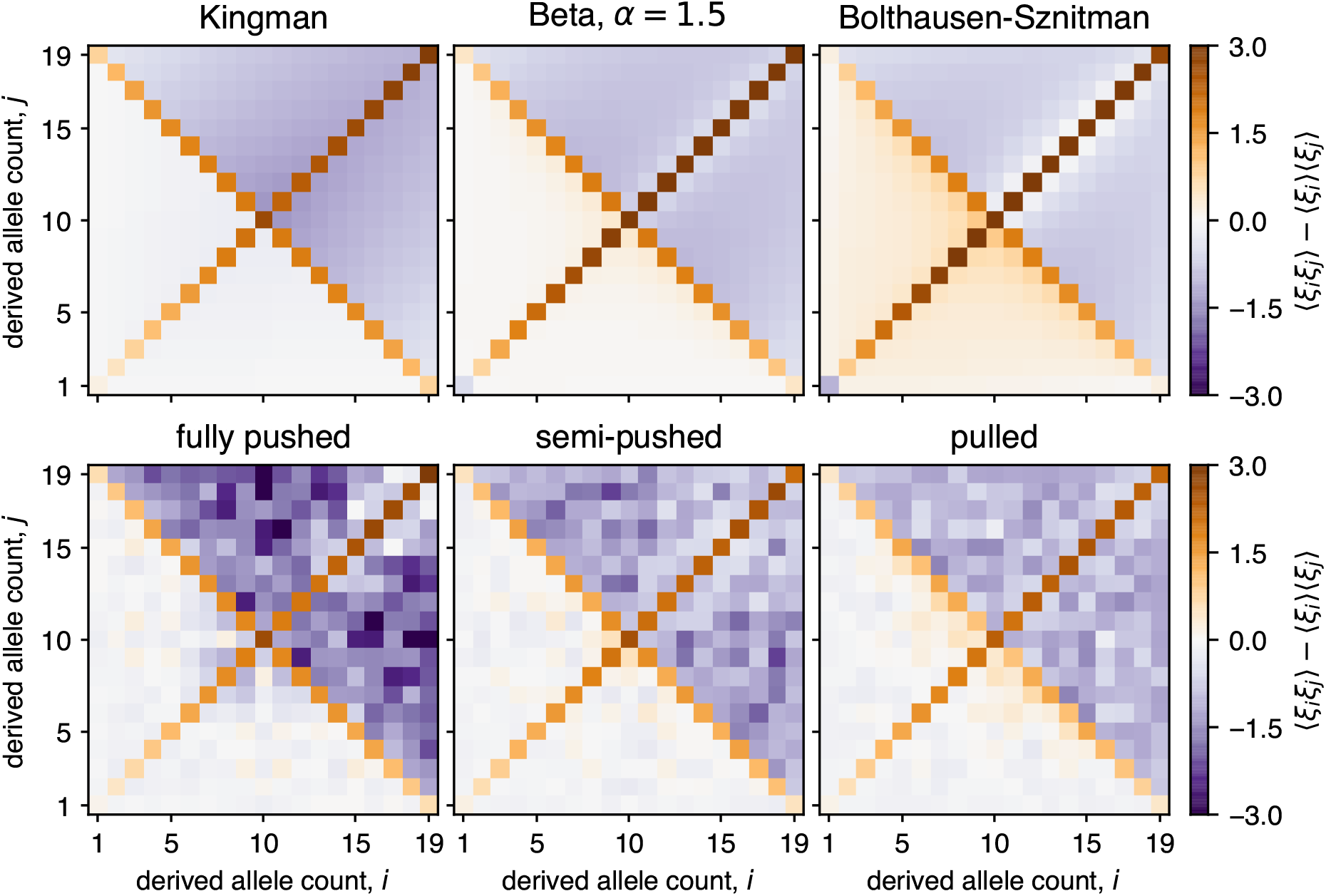
Comparison of two-site frequency spectra reveal signatures of multiple mergers in semi-pushed and pulled expansions. Matrices show the correlation function between tree branches subtending different number of leaves for both the expected coalescents (top) and expansion simulations (bottom) for each expansion regime. The averaged 2-SFS from simulations were generated using the same sampling procedure used for the SFS (SI, Sec. IV).

### IV. Simulations

In this section we explain the details of our expansion simulations and the data collection and processing pipelines.

#### Expansion simulations

We simulated the expansion of a population in a one-dimensional habitat modeled by an array of patches (demes), separated by a distance Δ*x*. Demes contain individuals, which are labeled using integers. We denote by *I_i_*(*t, x*) the label of the ith individual in deme *x*, with 1 ≤ *x* ≤ *L* and 1 ≤ *i* ≤ *N*. To allow for demes with less than *N* individuals, we use vacancies, which are labeled by *I^v^* = 0.

**Figure S2:**
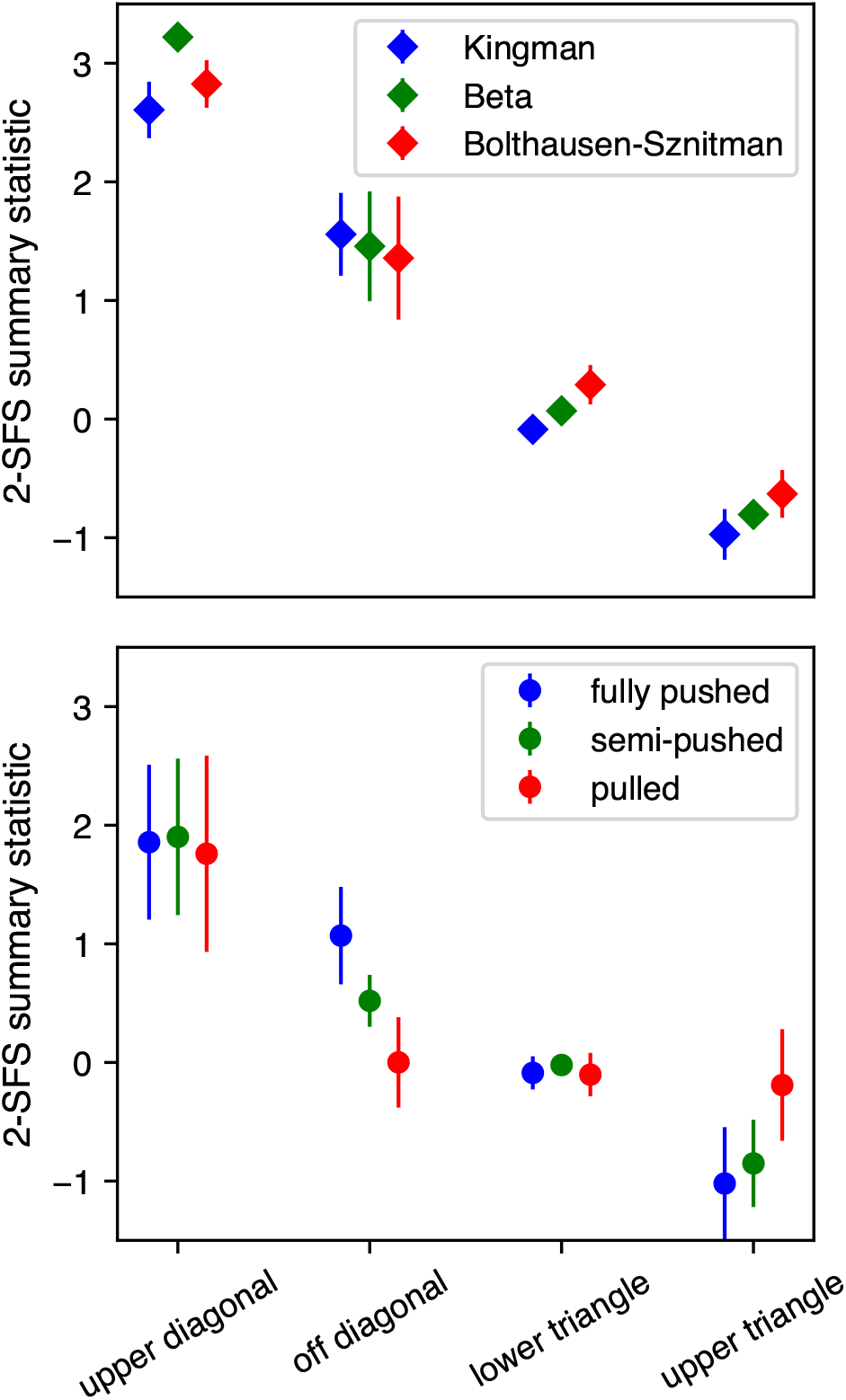
Summary statistics of 2-SFS show qualitative agreement with theoretical predictions. (Top) Mean values and standard deviation of entries in the 2-SFS for the Kingman (blue), Beta-with *β* = 1.5 (green), and Bolthausen-Sznitman (red) coalescents. For all three coalescents a sample size of *n* = 20 was used. The four bins are defined as follows: upper diagonal = {(*i,i*): ⌊*n*/2⌋ + 1 ≤ *i* ≤ *n* — 1}, off diagonal = {(*i,n — i*): 1 ≤ *i* < ⌊*n*/2⌋ or ⌊*n*/2⌋ < *i* < *n*}, lower triangle = {(*i,j*): *i* + *j* < n — 1,*i* = *j*}, upper triangle = {(*i,j*): *i* + *j > n* — 1,1 ≤ *i ≤ n* — 1,1 ≤ *j ≤ n* — 1,*i* = *j*}. (Bottom) Same as upper panel, but using 2-SFS from simulations of fully pushed (blue), semi-pushed (green) and pulled (red) expansions. All simulation parameters are identical to those for Fig. S1.

The population is initially localized on *L*/2 = 100 demes. Each deme is filled with *N* individually labeled members of the population. Individuals are labeled sequentially, starting with the first individual in the leftmost deme and moving to the right of the population. Thus,

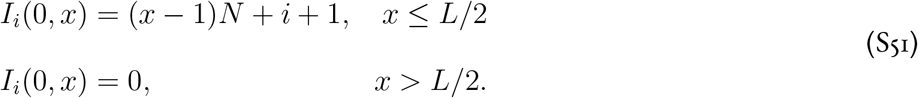

Each generation is updated in two steps. First, a migration step, in which demes are updated sequentially, starting from *x* =1. For each deme, the number of migrants exchanged with the next deme is drawn from a binomial distribution:

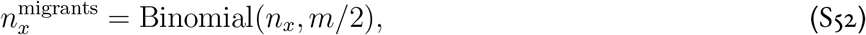

where *m* is the migration probability. To choose the migrants, the order of individuals in demes *x* and *x*+1 is randomized, and the first 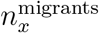 from the demes are exchanged.

Second, we perform a growth step. Following Ref. [2], the growth of the population was modeled by introducing a fitness difference between the vacancies and the actual species. Specifically, the fitness of the species was set to 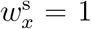 and the fitness of the vacancies was set to 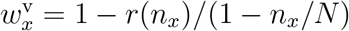, where

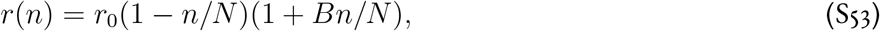

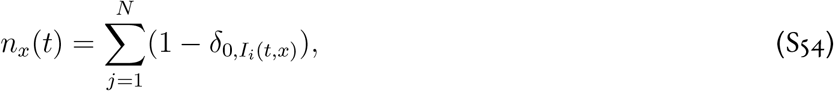

and *δ_lm_* is the Kronecker delta. The next generation is constructed by sampling, with replacement, a new set of labels *I_i_*(*t + 1,x*) from the set of previous labels {*I*_1_(*t, x*), *I*_2_(*t, x*),..., *I_N_* (*t,x*)} for each *i ≤ N*. The probability to sample the ancestor *I_i_*(*t, x*) is proportional to the ratio of *w_ix_* to the mean fitness of the population in the deme: 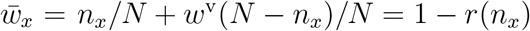.

#### Recording genealogies

The genealogy of the population is recorded in a custom tree class, in which all individuals in the simulation box are stored as nodes. Each node is assigned a unique parent node, and a set of child nodes, except for the most recent generation, which have no children—we will refer to these nodes as the leaves of the tree. The tree is initialized with one node, which is designated as the root of the tree, and is continuously updated as follows. At the start of the simulation, all individuals at the front are assigned as leaves with the root as their parent. As the population expands, many labels become extinct and the average clone size of the surviving labels grows. After a fixed number of generations Δ*t*, we relabel all individuals and add them as new nodes on the tree. Each individual is assigned as a child node to one of the leaves of the tree, which is designated as its parent according to the previous label of the child node. After every individual is assigned to the tree, the newly added nodes are designated as the new leaves of the tree. At the end of this process, we prune the tree by removing all nodes which have no leaves among their descendants. The process is repeated until either the whole population has one common ancestor or a maximum number of generations *T*_max_ for the simulation is reached.

### V. Data analysis

In this section we explain how we analyze the data from simulations to obtain the figures in the main text and the SI.

#### Estimating *τ_m_*

We used the following procedure to determine the spatial distribution of ancestors in Fig. 2 in the main text. We ran 1000 simulations of a fully pushed expansion, for which we estimated the coalescence time *T_c_* ≈ 10^3^, using the following parameters: *N* = 350, *B* = 10, *r*_0_ = 0.01, *m* = 0.4, *Δ*t** = 20. For each simulation we recorded the ancestry as described in Sec. IV. In order to determine the location of each ancestor from the population, we modified the label assignment algorithm by using the following equation:

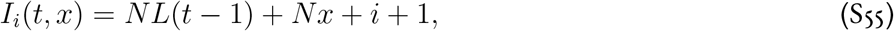

where *I_i_*(*t, x*) is the label of individual *i* from deme *x* in generation t. Using this equation, each label uniquely specifies the position of the individual.

Because fronts are stochastic it is difficult to compare ancestral distributions across simulations. To minimize the variance in the ancestral distribution due to variations in the final sampling location, we used the following procedure. We first determined the midpoint of the front, given by the deme closest to the mean position *x* along the front, weighted by the population size at *x*. Next, we determined the bulk and leading edge of the front, which we defined as least advanced location with population size below the carrying capacity and the most advanced location with a non-zero population size, respectively. Finally, the sampling location from the bulk and the front were chosen as the closest demes to the halfway distance between midpoint of the front, and the bulk edge and the front edge, respectively.

We then collect the labels of all individuals from the two sampling locations. Using the ancestral trees, we traced back the labels of the ancestors of all the sampled individuals. Finally, we recorded the locations of these ancestors by solving for *x* in Eq. (S55) and plotted the distribution of these locations across all simulations.

#### Sampling and analysis of SFS and 2-SFS

We used the following procedure to sample and analyze the SFS and 2-SFS from the ancestral trees in our simulations. We first subsampled a number of individuals *n* from close to the edge of the front in the final population. As discussed in the main text, far from the front the effects of spatial structure become important and our well-mixed approximation breaks down. Empirically, we observed that sampling individuals from the farthest advanced 20 demes minimized the effects of spatial structure on both the SFS and the clone size distributions shown in Fig. S4. The value of *n* was chosen small enough to allow for comparison with the exact predictions for the different colescent classes described below. Each ancestral tree was sampled independently 10 times in order to obtain better estimates for the averaged quantities we calculated.

The simulations used to generate the ancestral trees were performed choosing three values of *B* (10, 3.33, and 0, respectively) in Eq. (1) for each class of waves. Using Eq. (4) from the main text corresponding, the values of *α* for each of these expansions are approximately 1, 0.5, and 0, respectively. The coalescents for well-mixed populations with these descendant distributions are described by the Beta-coalescent from Eq.(S50) with the paramter *β* equal to 2, 1.5, and 1, respectively. To calculate the theoretical predictions for the SFS and 2-SFS we adapted a numerical implementation of the exact recurrence relations for the SFS and 2-SFS from Ref. [27], which was originally developed in Ref. [26].

### VI. Supplemental figures

**Figure S3:**
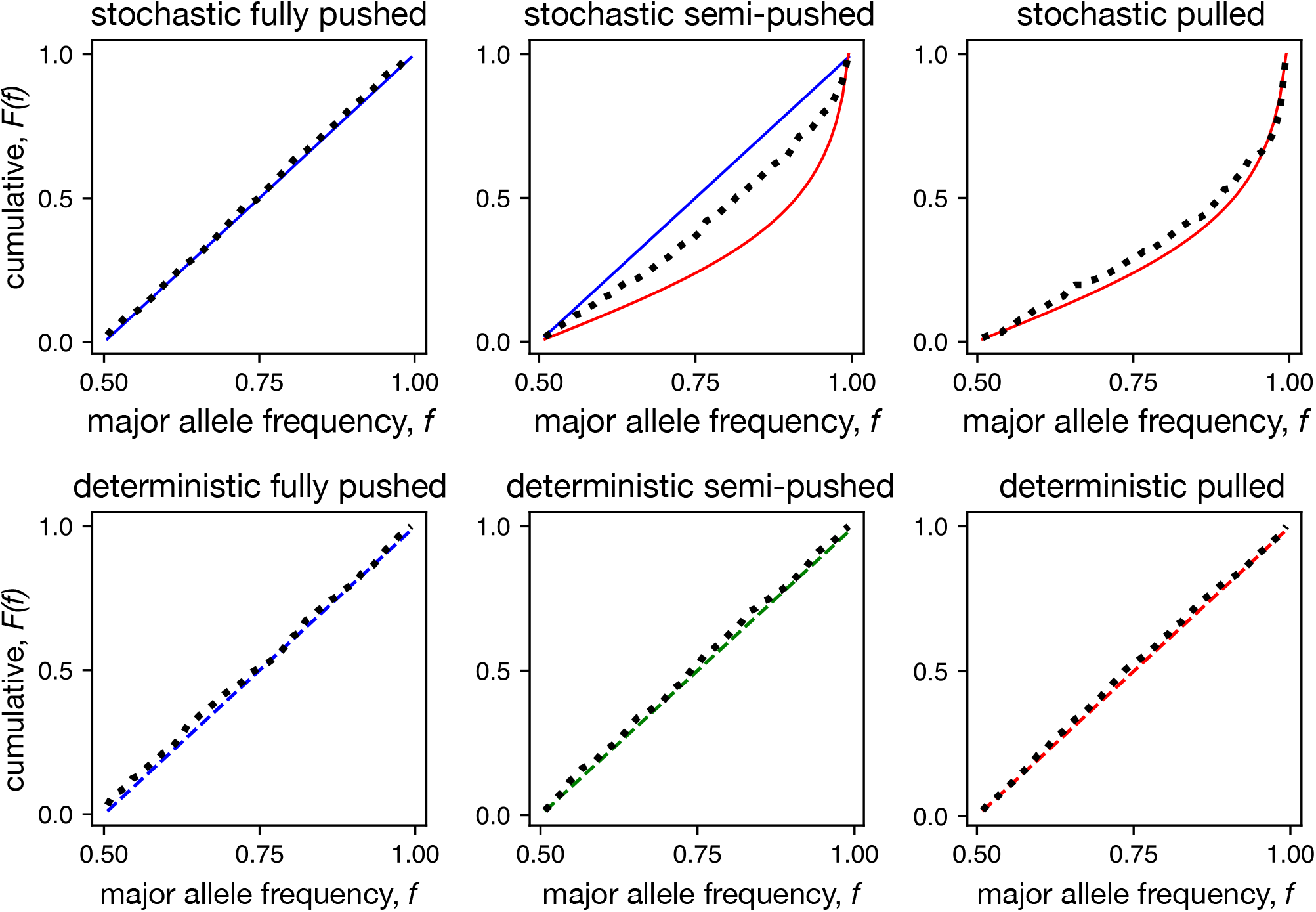
Allele frequency distributions quantitatively agree with theoretical predictions in both stochastic and deterministic regimes. Shows the same data as Fig. 5 in the main text as a cumulative distribution for better quantitative comparison between theoretical prediction and simulations. Simulations were carried out using the following parameters: *N* = 10^6^, *r*_0_ = 0.01, *m* = 0.4, *B* =10 (fully pushed), *B* = 3.33 (semi-pushed), and *B* = 0 (pulled). All simulations were started with equal frequency of the two alleles across the front. Distributions here and in Fig. 5 are shown after 3, 980, 000 (fully pushed), 1, 527, 315 (semi-pushed), and 98,827 (pulled) generations from the start for stochastic waves and after 4, 980, 000 (fully pushed), 1, 507, 719 (semi-pushed), and 531, 529 (pulled) generations for deterministic waves.

**Figure S4:**
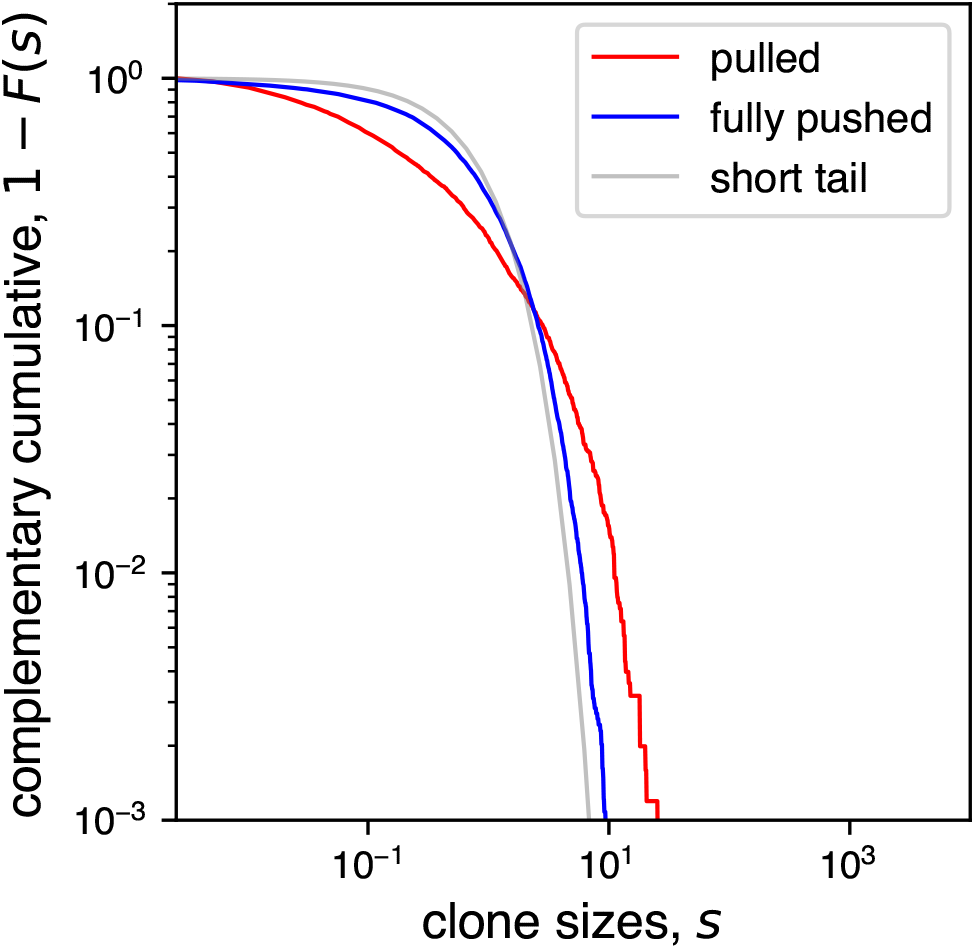
Pulled expansions have broader clone size distribution compared to fully pushed expansions. Left panel shows complementary cumulative distribution function of the normalized clone size *s* (where the normalization is with respect to the mean clone size) for fully pushed and pulled expansions. A total of 100 simulations were run without relabeling individuals and the sizes of distinct clones at the edge of the front were recorded every 500 generations. The front was defined as the first 25 demes starting from the most advanced occupied deme. The right panel shows the same data as cumulative distribution to emphasize the differences for small clone sizes. The growth function used is given by Eq. (1) with *B* =10 (fully pushed) and *B* = 0 (pulled) and all other parameters kept constant. The values of the other simulation parameters were *N* = 9600, *r*_0_ = 0.01, *m* = 0.4.

## Footnotes

1 Such expansions fall within the broader class of “pulled” expansions and we will usually refer to them by this term. Subsequent work rigorously proved that fitness waves are described by the Bolthausen-Sznitman coalescent [41], but no such proof exists for pulled spatial expansions, to our knowledge.

2 The fixation probability *u*(*ζ*) is always monotonic in *ζ* for pulled and semi-pushed expansions, and therefore *W_c_* ∝ *u*(*ζ_c_*). In fully pushed expansions, *u*(*ζ*) can have a maximum at *ζ < ζ_c_*, in which case there would be no cutoff in *P* (*W*). However, since fully pushed expansions are described by the Kingman coalescent, this does not change the conclusions of our argument.

3 Other summary statistics have also been used to describe the shape of genealogical trees. Perhaps the most popular of these is the total tree length, which determines the number of segregating sites in sequencing data. However, this metric is known to be very sensitive demographic expansions and is not a reliable indicator of coalescents with multiple mergers [52,53].

4 The critical value is determined from (4) by finding the value of *v/v_F_* for which *α* = 1.

5 This prediction assumes the population size is infinite, in which case P(f) widens in time and there is no strictly stationary distribution [61]. However, within the range of 1 /*N_e_* ≪ *f* ≪ 1 — 1 /*N_e_* we expect the allele frequency distribution to match the theoretical prediction, as we indeed see in simulations.

6 Mathematically, this property is referred to as exchangeability and is an underlying premise in coalescent theory.

7 This result applies to all Λ-coalescents, in which any number of lineages can merge at the same time, but merger events happen in succession. An even more general class, known as the Ξ-coalescent, allows for multiple simultaneous merger events as well. Such models have mainly been used to describe genealogies of diploid populations [19–21], but also populations under strong selection in the presence of recombination [22, 23].

## References

1. Donnelly, P. & Tavaré, S. Coalescents and genealogical structure under neutrality. Annu. Rev. Genet. 29, 401–421. ISSN: 00664197 (1995).

2. Li, H. & Durbin, R. Inference of human population history from individual whole-genome sequences. Nature 475, 493–496. ISSN: 1476-4687. https://doi.org/10.1038/nature10231 (July 2011).

3. Fujita, M. K., Leache, A. D., Burbrink, F. T., McGuire, J. A. & Moritz, C. Coalescent-based species delimitation in an integrative taxonomy. Trends in Ecology & Evolution 27, 480–488. ISSN: 0169-5347. http://www.sciencedirect.com/science/article/pii/S0169534712001000 (Sept. 2012).

4. Dayarian, A. & Shraiman, B. I. How to Infer Relative Fitness from a Sample of Genomic Sequences. Genetics 197, 913. http://www.genetics.org/content/197/3/913.abstract (July 2014).

5. Neher, R. A., Russell, C. A. & Shraiman, B. I. Predicting evolution from the shape of genealogical trees. Elife 3, 1–18. ISSN: 2050084X. arXiv: 1406.0789 (2014).

6. Kingman, J. F. C. The coalescent. Stochastic processes and their applications 13, 235–248 (1982).

7. Berestycki, N. Recent progress in coalescent theory. Ensaios Matematicos 16, 1–193 (2009).

8. Sella, G., Petrov, D. A., Przeworski, M. & Andolfatto, P. Pervasive Natural Selection in the Drosophila Genome? PLOS Genetics 5. Publisher: Public Library of Science, e1000495. https://doi.org/10.1371/journal.pgen.1000495 (June 2009).

9. Corbett-Detig, R. B., Hartl, D. L. & Sackton, T. B. Natural Selection Constrains Neutral Diversity across A Wide Range of Species. PLOS Biology 13. Publisher: Public Library of Science, e1002112. https://doi.org/10.1371/journal.pbio.1002112 (Apr. 2015).

10. Kern, A. D. & Hahn, M. W. The Neutral Theory in Light of Natural Selection. Molecular Biology and Evolution 35, 1366–1371. ISSN: 0737-4038. https://doi.org/10.1093/molbev/msy092 (2019) (May 2018).

11. Menardo, F., Gagneux, S. & Freund, F. Multiple merger genealogies in outbreaks of Mycobacterium tuberculosis. bioRxiv 12, 885723 (2019).

12. Hudson, R. R. & Kaplan, N. L. The coalescent process in models with selection and recombination. Genetics 120, 831. http://www.genetics.org/content/120/3/831.abstract (Nov. 1988).

13. Nei, M. & Takahata, N. Effective population size, genetic diversity, and coalescence time in subdivided populations. Journal of Molecular Evolution 37. Publisher: Springer, 240–244. ISSN: 0022-2844 (1993).

14. Wakeley, J. Distinguishing migration from isolation using the variance of pairwise differences. Theoretical population biology 49. Publisher: Elsevier, 369–386. ISSN: 0040-5809 (1996).

15. Nordborg, M. Structured Coalescent Processes on Different Time Scales. Genetics 146, 1501. http://www.genetics.org/content/146/4/1501.abstract (Aug. 1997).

16. Charlesworth, B., Charlesworth, D. & Barton, N. H. The effects of genetic and geographic structure on neutral variation. Annual Review of Ecology, Evolution, and Systematics 34, 99–125 (2003).

17. Wakeley, J. & Aliacar, N. Gene Genealogies in a Metapopulation. Genetics 159, 893. http://www.genetics.org/content/159/2/893.abstract (Oct. 2001).

18. Sargsyan, O. & Wakeley, J. A coalescent process with simultaneous multiple mergers for approximating the gene genealogies of many marine organisms. Theoretical Population Biology 74, 104–114. ISSN: 0040-5809. http://www.sciencedirect.com/science/article/pii/S0040580908000580 (Aug. 2008).

19. Rodelsperger, C. et al. Characterization of Genetic Diversity in the Nematode iem¿Pristionchus pacificusi/em¿ from Population-Scale Resequencing Data. Genetics 196, 1153. http://www.genetics.org/content/196/4/1153.abstract (Apr. 2014).

20. Schweinsberg, J. Coalescents with simultaneous multiple collisions. Electronic Journal of Probability 5 (2000).

21. Eldon, B. & Wakeley, J. Coalescent processes when the distribution of offspring number among individuals is highly skewed. Genetics 172, 2621–2633 (2006).

22. Pitman, J. Coalescents with multiple collisions. Annals of Probability 27. Publisher: JSTOR, 1870–1902. ISSN: 0091-1798 (Oct. 1999).

23. Ramachandran, S. et al. Support from the relationship of genetic and geographic distance in human populations for a serial founder effect originating in Africa. Proceedings of the National Academy of Sciences of the United States of America 102, 15942–15947 (2005).

24. Pierce, A. A. et al. Serial founder effects and genetic differentiation during worldwide range expansion of monarch butterflies. Proceedings of the Royal Society B: Biological Sciences 281. Publisher: Royal Society, 20142230. https://doi.org/10.1098/rspb.2014.2230 (2020) (Dec. 2014).

25. Britton, J. R. & Gozlan, R. E. How many founders for a biological invasion? Predicting introduction outcomes from propagule pressure. Ecology 94. Publisher: John Wiley & Sons, Ltd, 2558–2566. ISSN: 0012-9658. https://doi.org/10.1890/13-0527.1 (2020) (Nov. 2013).

26. Phillips, B. L., Brown, G. P., Webb, J. K. & Shine, R. Invasion and the evolution of speed in toads. Nature 439. Publisher: Nature Publishing Group, 803–803. ISSN: 1476-4687 (2006).

27. Hellberg, M. E., Balch, D. P. & Roy, K. Climate-Driven Range Expansion and Morphological Evolution in a Marine Gastropod. Science 292, 1707. http://science.sciencemag.org/content/292/5522/1707.abstract (June 2001).

28. Hallatschek, O., Hersen, P., Ramanathan, S. & Nelson, D. R. Genetic drift at expanding frontiers promotes gene segregation. Proceedings of the National Academy of Sciences 104, 19926–19930 (2007).

29. Cremer, J. et al. Chemotaxis as a navigation strategy to boost range expansion. Nature 575, 658–663. ISSN: 1476-4687. https://doi.org/10.1038/s41586-019-1733-y (Nov. 2019).

30. Gerlee, P. & Nelander, S. The impact of phenotypic switching on glioblastoma growth and invasion. PLoS Computational Biology 8, e1002556 (2012).

31. Sottoriva, A. et al. A Big Bang model of human colorectal tumor growth. Nature Genetics 47, 209–216. ISSN: 1546-1718. https://doi.org/10.1038/ng.3214 (Mar. 2015).

32. Slatkin, M. & Excoffier, L. Serial founder effects during range expansion: A spatial analog of genetic drift. Genetics 191, 171–181. ISSN: 00166731 (2012).

33. DeGiorgio, M., Jakobsson, M. & Rosenberg, N. A. Explaining worldwide patterns of human genetic variation using a coalescent-based serial founder model of migration outward from Africa. Proceedings of the National Academy of Sciences 106, 16057. http://www.pnas.org/content/106/38/16057.abstract (Sept. 2009).

34. Brunet, E., Derrida, B., Mueller, A. H. & Munier, S. Effect of selection on ancestry: an exactly soluble case and its phenomenological generalization. Physical Review E 76, 041104 (2007).

35. Excoffier, L., Foll, M. & Petit, R. J. Genetic Consequences of Range Expansions. Annual Review of Ecology, Evolution, and Systematics 40, 481–501. http://www.annualreviews.org/doi/abs/10.1146/annurev.ecolsys.39.110707.173414 (Dec. 2009).

36. DeGiorgio, M., Degnan, J. H. & Rosenberg, N. A. Coalescence-Time Distributions in a Serial Founder Model of Human Evolutionary History. Genetics 189, 579. http://www.genetics.org/content/189/2/579.abstract (Oct. 2011).

37. Etheridge, A. & Penington, S. Genealogies in bistable waves. arXiv preprint 9, 2009.03841 (2020).

38. Tsimring, L. S., Levine, H. & Kessler, D. A. RNA virus evolution via a fitness-space model. Physical review letters 76, 4440 (1996).

39. Rouzine, I. M., Wakeley, J. & Coffin, J. M. The solitary wave of asexual evolution. Proceedings of the National Academy of Sciences 100, 587–592 (2003).

40. Hallatschek, O. The noisy edge of traveling waves. Proceedings of the National Academy of Sciences 108, 1783–1787 (2011).

41. Schweinsberg, J. Rigorous results for a population model with selection II: genealogy of the population. Electron. J. Probab. 22, 54 pp. https://doi.org/10.1214/17-EJP58 (2017).

42. Bolthausen, E. & Sznitman, A.-S. On Ruelle’s Probability Cascades and an Abstract Cavity Method. Communications in Mathematical Physics 197, 247–276. ISSN: 1432-0916. https://doi.org/10.1007/s002200050450 (Oct. 1998).

43. Neher, R. A. & Hallatschek, O. Genealogies of rapidly adapting populations. Proceedings of the National Academy of Sciences 110, 437–442 (2013).

44. Birzu, G., Hallatschek, O. & Korolev, K. S. Fluctuations uncover a distinct class of traveling waves. Proc. Natl. Acad. Sci. 115, E3645–E3654. https://doi.org/10.1073/pnas.1715737115 (Apr. 2018).

45. Birzu, G., Matin, S., Hallatschek, O. & Korolev, K. S. Genetic drift in range expansions is very sensitive to density dependence in dispersal and growth. Ecology Letters 22, 1817–1827. ISSN: 1461-023X. https://doi.org/10.1111/ele.13364 (2019) (Nov. 2019).

46. Sagitov, S. The general coalescent with asynchronous mergers of ancestral lines. Journal of Applied Probability 36, 1116–i125 (1999).

47. Schweinsberg, J. Coalescent processes obtained from supercritical Galton-Watson processes. Stochastic processes and their Applications 106, 107–139 (2003).

48. Kimura, M. & Weiss, G. H. The stepping stone model of population structure and the decrease of genetic correlation with distance. Genetics 49, 561. http://www.genetics.org/content/49/4/561.abstract (Apr. 1964).

49. Dennis, B. & Taper, M. L. Density Dependence in Time Series Observations of Natural Populations: Estimation and Testing. Ecological Monographs 64. Publisher: John Wiley & Sons, Ltd, 205–224. ISSN: 0012-9615. https://doi.org/10.2307/2937041 (2020) (Feb. 1994).

50. May, R. & McLean, A. R. Theoretical ecology:principles and applications ISBN: 0-19-920999-5 (Oxford University Press on Demand, 2007).

51. Hallatschek, O. & Nelson, D. R. Gene surfing in expanding populations. Theoretical Population Biology 73, 158–170 (2008).

52. Tajima, F. The effect of change in population size on DNA polymorphism. Genetics 123, 597. http://www.genetics.org/content/123/3/597.abstract (Nov. 1989).

53. Rice, D. P., Novembre, J. & Desai, M. M. Distinguishing multiple-merger from Kingman coalescence using two-site frequency spectra. bioRxivpreprint 11, 461517 (2018).

54. Hudson, R. R. Two-Locus Sampling Distributions and Their Application. Genetics 159, 1805. http://www.genetics.org/content/159/4/1805.abstract (Dec. 2001).

55. Ferretti, L. et al. The neutral frequency spectrum of linked sites. Theoretical Population Biology 123, 70–79. ISSN: 0040-5809. http://www.sciencedirect.com/science/article/pii/S0040580917301399 (Sept. 2018).

56. Fu, Y. Statistical Properties of Segregating Sites. Theoretical Population Biology 48, 172–197. ISSN: 0040-5809. http://www.sciencedirect.com/science/article/pii/S0040580985710258 (Oct. 1995).

57. Fisher, R. A. The Genetical Theory of Natural Selection (Oxford University Press, Oxford, United Kingdom, 1999).

58. Barton, N. H. & Etheridge, A. M. The relation between reproductive value and genetic contribution. Genetics 188, 953–973 (2011).

59. Gandhi, S. R., Korolev, K. S. & Gore, J. Cooperation mitigates diversity loss in a spatially expanding microbial population. Proceedings of the National Academy of Sciences 116, 23582–23587 (2019).

60. Mohle, M. & Sagitov, S. A classification of coalescent processes for haploid exchangeable population models. The Annals of Probability 29. Publisher: Institute of Mathematical Statistics, 1547–1562. ISSN: 0091-1798 (2001).

61. Hallatschek, O. Selection-like biases emerge in population models with recurrent jackpot events. Genetics 210, 1053–1073 (2018).

62. Donnelly, P. & Kurtz, T. G. Genealogical Processes for Fleming-Viot Models with Selection and Recombination. The Annals of Applied Probability 9. Publisher: Institute of Mathematical Statistics, 1091–1148. ISSN: 10505164. http://www.jstor.org/stable/2667143 (2020) (1999).

63. Kimura, M. Solution of a process of random genetic drift with a continuous model. Proceedings of the National Academy of Sciences 41, 144. http://www.pnas.org/content/41/3/144.abstract (Mar. 1955).

64. Austerlitz, F., Jung-Muller, B., Godelle, B. & Gouyon, P.-H. Evolution of coalescence times, genetic diversity and structure during colonization. Theoretical Population Biology 51. Publisher: Elsevier, 148–164. ISSN: 0040-5809 (1997).

65. Roques, L., Garnier, J., Hamel, F. & Klein, E. K. Allee effect promotes diversity in traveling waves of colonization. Proceedings of the National Academy of Sciences 109, 8828–8833 (2012).

66. Fierer, N., Nemergut, D., Knight, R. & Craine, J. M. Changes through time: integrating microorganisms into the study of succession. Research in Microbiology 161, 635–642. ISSN: 0923-2508. http://www.sciencedirect.com/science/article/pii/S0923250810001385 (Oct. 2010).

67. Challagundla, L. et al. Range Expansion and the Origin of USA300 North American Epidemic Methicillin-Resistant Staphylococcus aureus. mBio 9 (ed Barbour, A. G.) eprint: https://mbio.asm.org/content/9/1/e02016-17.full.pdf. https://mbio.asm.org/content/9/1/e02016-17 (2018).

68. Phillips, B. L., Brown, G. P., Greenlees, M., Webb, J. K. & Shine, R. Rapid expansion of the cane toad (Bufo mari-nus) invasion front in tropical Australia. Austral Ecology 32, 169–176 (2007).

69. Gray, M. E., Sappington, T. W., Miller, N. J., Moeser, J. & Bohn, M. O. Adaptation and invasiveness of western corn rootworm: intensifying research on a worsening pest. Annual review of entomology 54, 303–321 (2009).

70. Pateman, R. M., Hill, J. K., Roy, D. B., Fox, R. & Thomas, C. D. Temperature-dependent alterations in host use drive rapid range expansion in a butterfly. Science 336, 1028–1030 (2012).

71. Reiter, M., Rulands, S. & Frey, E. Range Expansion of Heterogeneous Populations. Physical Review Letters 112. Publisher: American Physical Society, 148103. https://link.aps.org/doi/10.1103/PhysRevLett.112.148103 (Apr. 2014).

72. Marculis, N. G., Lui, R. & Lewis, M. A. Neutral genetic patterns for expanding populations with nonoverlapping generations. Bulletin of Mathematical Biology 79, 828–852 (2017).

73. Klopfstein, S., Currat, M. & Excoffier, L. The Fate of Mutations Surfing on the Wave of a Range Expansion. Molecular Biology and Evolution 23, 482–490. ISSN: 0737-4038. https://doi.org/10.1093/molbev/msj057 (2020) (Mar. 2006).

74. Kessler, D. A., Ner, Z. & Sander, L. M. Front propagation: precursors, cutoffs, and structural stability. Physical ReviewE 58, 107 (1998).

75. Desai, M. M., Walczak, A. M. & Fisher, D. S. Genetic diversity and the structure of genealogies in rapidly adapting populations. Genetics 193, 565–585 (2013).

76. Good, B. H., Walczak, A. M., Neher, R. A. & Desai, M. M. Genetic Diversity in the Interference Selection Limit. PLOS Genetics 10, e1004222. https://doi.org/10.1371/journal.pgen.1004222 (Mar. 2014).

77. Schrider, D. R., Shanku, A. G. & Kern, A. D. Effects of Linked Selective Sweeps on Demographic Inference and Model Selection. Genetics 204, 1207. http://www.genetics.org/content/204/3/1207.abstract (Nov. 2016).

## References

1. Roques, L., Garnier, J., Hamel, F. & Klein, E. K. Allee effect promotes diversity in traveling waves of colonization. Proceedings of the National Academy of Sciences 109, 8828–8833 (2012).

2. Hallatschek, O. & Nelson, D. R. Gene surfing in expanding populations. Theoretical Population Biology 73, 158–170 (2008).

3. Birzu, G., Hallatschek, O. & Korolev, K. S. Fluctuations uncover a distinct class of traveling waves. Proc. Natl. Acad. Sci. 115, E3645–E3654. https://doi.org/10.1073/pnas.1715737115 (Apr. 2018).

4. Hallatschek, O. Selection-like biases emerge in population models with recurrent jackpot events. Genetics 210, 1053–1073 (2018).

5. Berestycki, N. Recent progress in coalescent theory. Ensaios Matematicos 16, 1–193 (2009).

6. Kimura, M. Solution of a process of random genetic drift with a continuous model. Proceedings of the National Academy of Sciences 41, 144. http://www.pnas.org/content/41/3/144.abstract (Mar. 1955).

7. Watson, H. W. & Galton, F. On the probability of the extinction of families. The Journal of the Anthropological Institute of Great Britain and Ireland 4, 138–144 (1875).

8. Harris, T. E. The theory of branching processes (Courier Corporation, 2002).

9. Durrett, R. Probability: theory and examples (Cambridge university press, 2010).

10. Harris, T. E. Some mathematical models for branching processes tech. rep. (RAND CORP SANTA MONICA CA, 1950).

11. Otter, R. The multiplicative process. The Annals of Mathematical Statistics, 206–224 (1949).

12. Kolmogorov, A. N. On the solution of a biological problem. Proceedings of Tomsk University 2, 7–12 (1938).

13. Zolotarev, V. M. More exact statements of several theorems in the theory of branching processes. Theory of Probability & Its Applications 2, 245–253 (1957).

14. Goh, K.-I., Lee, D.-S., Kahng, B. & Kim, D. Sandpile on Scale-Free Networks. Physical Review Letters 91. Publisher: American Physical Society, 148701. https://link.aps.org/doi/10.1103/PhysRevLett.91.148701 (Oct. 1, 2003).

15. Gleeson, J. P., Lee, W. T., Ward, J. A. & O’Sullivan, K. P. Competition-induced criticality in a model of meme popularity. Phys. Rev. Lett. 112, 048701 (2014).

16. Foucart, C., Henard, O., et al. Stable continuous-state branching processes with immigration and Beta-Fleming-Viot processes with immigration. Electronic Journal of Probability 18 (2013).

17. Li, Z. Continuous-state branching processes. arXiv preprint arXiv:1202.3223 (2012).

18. Kyprianou, A. E. & Pardo, J.-C. Continuous-state branching processes and self-similarity. Journal of Applied Probability 45, 1140–1160 (2008).

19. Mohle, M. & Sagitov, S. Coalescent patterns in diploid exchangeable population models. Journal of Mathematical Biology 47, 337–352. ISSN: 1432-1416. https://doi.org/10.1007/s00285-003-0218-6 (Sept. 1, 2003).

20. Birkner, M., Blath, J. & Eldon, B. An Ancestral Recombination Graph for Diploid Populations with Skewed Offspring Distribution. Genetics 193, 255. http://www.genetics.org/content/193/1/255.abstract (Jan. 1, 2013).

21. Birkner, M., Liu, H., Sturm, A., et al. Coalescent results for diploid exchangeable population models. Electronic Journal of Probability 23 (2018).

22. Durrett, R. & Schweinsberg, J. Approximating selective sweeps. Theoretical Population Biology 66, 129–138. ISSN: 0040-5809. http://www.sciencedirect.com/science/article/pii/S0040580904000607 (Sept. 1, 2004).

23. Durrett, R. & Schweinsberg, J. A coalescent model for the effect of advantageous mutations on the genealogy of a population. Stochastic Processes and their Applications 115, 1628–1657. ISSN: 0304-4149. http://www.sciencedirect.com/science/article/pii/S0304414905000608 (Oct. 1, 2005).

24. Desai, M. M., Walczak, A. M. & Fisher, D. S. Genetic diversity and the structure of genealogies in rapidly adapting populations. Genetics 193, 565–585 (2013).

25. Neher, R. A. & Hallatschek, O. Genealogies of rapidly adapting populations. Proceedings of the National Academy of Sciences 110, 437–442 (2013).

26. Birkner, M., Blath, J. & Eldon, B. Statistical Properties of the Site-Frequency Spectrum Associated with Lambda Coalescents. Genetics 195, 1037. http://www.genetics.org/content/195/3/1037.abstract (Nov. 2013).

27. Rice, D. P., Novembre, J. & Desai, M. M. Distinguishing multiple-merger from Kingman coalescence using two-site frequency spectra. bioRxivpreprint 11, 461517 (2018).

